# *De novo* distillation of thermodynamic affinity from deep learning regulatory sequence models of *in vivo* protein-DNA binding

**DOI:** 10.1101/2023.05.11.540401

**Authors:** Amr M. Alexandari, Connor A. Horton, Avanti Shrikumar, Nilay Shah, Eileen Li, Melanie Weilert, Miles A. Pufall, Julia Zeitlinger, Polly M. Fordyce, Anshul Kundaje

**Affiliations:** Department of Computer Science, Stanford University, Stanford, CA 94305; Department of Genetics, Stanford University, Stanford, CA 94305; Department of Earth System Science, Stanford University, Stanford, CA 94305; Stowers Institute for Medical Research, Kansas City, MO, USA; Department of Biochemistry, Carver College of Medicine, University of Iowa, Iowa City, Iowa 52242, USA; The University of Kansas Medical Center, Kansas City, KS, USA; Department of Bioengineering, Stanford University, Stanford, CA 94305; ChEM-H Institute, Stanford University, Stanford, CA 94305; Chan Zuckerberg Biohub, San Francisco, CA 94110

**Keywords:** Transcription factors, high-throughput binding assays, deep learning, protein-DNA binding, gene regulation, binding affinity

## Abstract

Transcription factors (TF) are proteins that bind DNA in a sequence-specific manner to regulate gene transcription. Despite their unique intrinsic sequence preferences, *in vivo* genomic occupancy profiles of TFs differ across cellular contexts. Hence, deciphering the sequence determinants of TF binding, both intrinsic and context-specific, is essential to understand gene regulation and the impact of regulatory, non-coding genetic variation. Biophysical models trained on *in vitro* TF binding assays can estimate intrinsic affinity landscapes and predict occupancy based on TF concentration and affinity. However, these models cannot adequately explain context-specific, *in vivo* binding profiles. Conversely, deep learning models, trained on *in vivo* TF binding assays, effectively predict and explain genomic occupancy profiles as a function of complex regulatory sequence syntax, albeit without a clear biophysical interpretation. To reconcile these complementary models of *in vitro* and *in vivo* TF binding, we developed Affinity Distillation (AD), a method that extracts thermodynamic affinities *de-novo* from deep learning models of TF chromatin immunoprecipitation (ChIP) experiments by marginalizing away the influence of genomic sequence context. Applied to neural networks modeling diverse classes of yeast and mammalian TFs, AD predicts energetic impacts of sequence variation within and surrounding motifs on TF binding as measured by diverse *in vitro* assays with superior dynamic range and accuracy compared to motif-based methods. Furthermore, AD can accurately discern affinities of TF paralogs. Our results highlight thermodynamic affinity as a key determinant of *in vivo* binding, suggest that deep learning models of *in vivo* binding implicitly learn high-resolution affinity landscapes, and show that these affinities can be successfully distilled using AD. This new biophysical interpretation of deep learning models enables high-throughput *in silico* experiments to explore the influence of sequence context and variation on both intrinsic affinity and *in vivo* occupancy.

## Background

Transcription factors bind DNA in a sequence-specific manner to regulate gene expression and thereby maintain cellular homeostasis and regulate organismal development. As TFs exert control over cellular differentiation and immune responses, they are implicated in cardiovascular diseases, inflammatory disorders, and many cancers and represent potential drug targets (1–4). Further, complex trait-associated variants are enriched in regulatory DNA and often disrupt TF binding sites(5–7). Comprehensive characterization of TF binding is essential to understand gene regulation, predict and interpret disease-associated non-coding variation, design novel regulatory circuits in synthetic biology and rationally design transgenes for biotechnology and gene therapy (8–10). Consequently, a quantitative and predictive understanding of TF binding has been a longstanding goal in biology (11).

Thermodynamic approaches have been successfully applied to model and quantitatively predict interactions between individual TFs and naked DNA sequences *in vitro* (12) and to map comprehensive binding affinity landscapes (13). In a simple two-state model, the probability of binding follows a logarithmic relationship with TF concentration: TF binding increases linearly at concentrations below 50% occupancy (at which the concentration = *K*_d_) and then asymptotes once available sites have been saturated. A key strength of this model is its generalizability: in principle, we can quantitatively predict binding to any pool of DNA sequences as a function of TF concentrations as long as we know the affinity for each. In practice, however, the possible DNA sequence space is too large to directly measure *K*_d_s for every possible sequence of interest. Instead, researchers have developed models that can learn affinities from large experimental datasets that measure TF binding to many sequences. The most widely-used model is the position weight matrix (PWM), which is a linear mononucleotide model and an early application of the perceptron algorithm (14). PWMs represent binding specificity as the enrichment of a particular nucleotide at a given position within a TF target ‘motifs’ (15,16). These motifs can then be used to scan arbitrary sequences of interest and identify TF-specific binding sites (11,17,18).

In recent decades, a wide variety of technologies have been developed to profile TF specificity *in vitro* and create vast TF-specific motif catalogs. Universal PBMs (19) measure TF binding to all possible *k*– mers, genomic-context PBMs (20) measure TF binding to sequences of putative genomic binding sites, SELEX-seq (21) utilizes *in vitro* selection to explore DNA binding preferences across millions of sequences, and MITOMI (22) provides high-resolution affinity measurements of TF-DNA interactions. While motifs derived from the same data typically explain the majority of observed *in vitro* binding, a surprising amount of variance often remains unexplained, particularly when motifs are embedded within longer and more complex sequences (23,24). The presence of different sequence contexts, even when these contexts are not predicted to have any binding by traditional PWM models, can have strong impacts (positive or negative) on actual binding energies (20,25–33).

The performance of motif-based models suffers further when attempting to predict genome-wide TF binding to chromatinized DNA in cells (Supplement **Fig. S1**). New and powerful assays capable of quantifying genome-wide TF binding in cells (*e.g.* ChIP-seq (34,35), ChIP-exo (36), ChIP-nexus (37), CUT&RUN (38,39), *etc.*) have yielded >500 and >2600 publicly available immunoprecipitation datasets for the Yeast Epigenome (40) and human/mouse ENCODE (41) initiatives. While high-affinity *in vitro*– derived motifs for a given TF are typically among the most enriched sequences in a ChIP-seq dataset, many genomic loci predicted to be bound are not occupied, and many TFs bind loci lacking apparent motifs (42). This discrepancy could reflect an increased importance of additional extrinsic factors beyond TF-DNA affinity in specifying proper localization in cells (*e.g.* cooperative and competitive interactions between TFs, local chromatin state, and three-dimensional chromatin architecture (11,43)) or could simply reflect the fact that motif-based additive models that do not capture impacts of sequence context necessary to accurately model specificity (24). The current failure of *in vitro*-derived specificity models to accurately predict TF occupancy *in vivo* is often cited as evidence that the thermodynamics of TF/DNA interactions are not the primary determinants of binding in cells. Here, we ask the question in an alternate way: to what extent do state-of-the-art models learn the thermodynamics of TF/DNA interactions when predicting *in vivo* occupancies?

Convolutional neural networks (CNNs) are a class of deep learning models that can map complex inputs (e.g. DNA sequences) to associated outputs (e.g. measures of TF binding occupancy) by learning complex, non-linear functions composed of hierarchical banks of *de novo* pattern detectors. A CNN with a single pattern detector (convolutional filter) can learn the same scoring function as a simple PWM. However, by using multiple hierarchical banks of multiple convolutional filters, CNNs can easily encode arbitrarily complex higher-order patterns of motifs and their dependencies enabling them to learn cis-regulatory sequence syntax without explicit knowledge or specification of these features. Hence, CNNs (e.g. DeepBind (44), DeepSEA (45), BPNet (46) and Enformer(47)) have been recently used to learn predictive sequence models of *in vivo* transcription factor binding. These models have proven useful to predict effects of genetic variants on context-specific TF binding(45,47). Further, model interpretation frameworks have been developed to explain the predictions of these models thereby enhancing our understanding of TF target site sequence syntax, impacts of cooperation and competition between TFs, effects of nucleosomes, and chromatin-mediated cooperativity (46). However, these models typically cannot extract quantitative information about binding affinities or how they may influence occupancy. We therefore still lack the pivotal link that connects state-of-the-art models with a scalable and predictive biophysical understanding of TF binding.

To address this gap, we introduce Affinity Distillation (AD), which leverages a recently developed and highly accurate convolutional neural network model (BPNet) and a marginalization technique to allow high-throughput determination of thermodynamic affinities from abundant and physiologically relevant *in vivo* occupancy data (**Fig. 1A**). AD systematically injects sequences of interest within hundreds of randomized sequence contexts and then uses trained BPNet models to predict summed binding counts, thereby isolating the energetic impacts of a sequence of interest. This *in silico* marginalization at scale results in scores that are strongly linearly correlated with *in vitro*-measured relative binding energies (ΔΔGs) after training on a wide variety of data types (PB-exo, ChIP-seq, and ChIP-nexus) from both yeast and human cells. In all cases, systematic comparisons between AD and other best-in-class predictive approaches establish that AD outperforms existing motif-based and NN-based methods. AD also successfully predicts subtle differences in binding between closely related paralogs that share similar consensus motifs, enabling differential analyses not previously possible using motifs. Finally, AD can adjust predictions non-additively, for instance to properly model non-specific binding, which additive models cannot. AD is therefore able to overcome previous limitations of motif-based methods and, by making accurate predictions that minimize the effect of genomic sequence context, decipher the thermodynamic contributions of sequence affinities.

**Figure 1.**
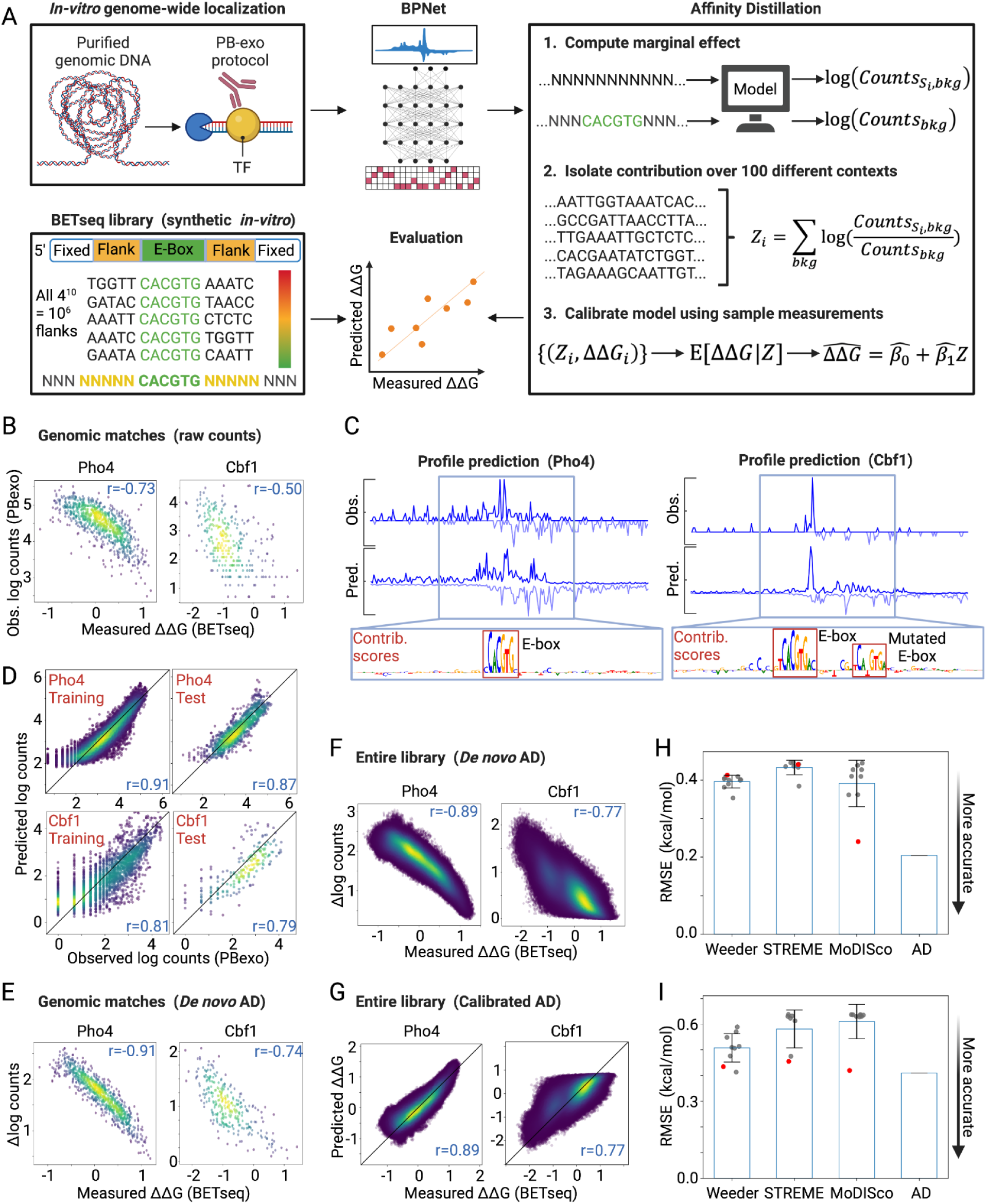
**Affinity Distillation extracts affinities from PB-exo models in yeast.** (A) Pipeline overview: BPNet models are trained on PB-exo data for Pho4 and Cbf1, Affinity Distillation (AD) extracts affinities, and AD outputs are compared with BETseq-measured ΔΔGs for Pho4 and Cbf1 binding to all 1,048,576 NNNNNCACGTGNNNNN sequences. (B) Observed and predicted log-transformed read counts for Pho4 (top) and Cbf1 (bottom) for the training (left) and held-out test chromosomes (right). (C) Observed (Obs) and predicted (Pred) PB-exo profile for sample regions located on held-out test chromosomes; contribution scores highlight known CACGTG consensus sites within bound regions. (D) Measured log-transformed PB-exo read counts vs measured ΔΔGs for NNNNNCACGTGNNNNN sequences present in both PB-exo and BET-seq experiments (884 and 294 sequences for Pho4 and Cbf1). (E) AD-predicted marginalization scores vs measured ΔΔGs for NNNNNCACGTGNNNNN sequences present in both PB-exo and BET-seq experiments. (F) AD-predicted marginalization scores vs measured ΔΔGs for all 1,048,576 NNNNNCACGTGNNNNN sequences in BET-seq experiments. (G) Calibrated AD predictions vs measured ΔΔGs for all 1,048,576 NNNNNCACGTGNNNNN sequences in BET-seq experiments. (H) Post-calibration RMSEs of predictions vs observations for Pho4. Gray markers indicate individual motif performances; red markers indicate the performance of the top output of each algorithm; blue bars indicate mean values; error bars indicate standard deviation. (I) Same as H for Cbf1.

## Results

### Affinity Distillation uses *in silico* marginalization to isolate the thermodynamic contribution of an arbitrary DNA sequence

BPNet takes one-hot encoded DNA sequences and predicts a decomposed binding profile: the total signal (read counts) and the profile shape (base-resolution distribution of reads). To estimate affinities for a sequence of interest, we trained BPNet models on 1 kb regions surrounding reproducible peaks within each dataset (**Fig. 1A**). In each case, we tuned hyperparameters and evaluated performance using different sets of genomic regions in distinct chromosomes (see Methods). After training, we interpreted the models using DeepSHAP, which decomposes model output predictions into contribution scores from individual nucleotides. We evaluated the resulting models by: (1) computing Pearson correlations and root mean squared error between predicted and observed read counts across regions from test chromosomes, (2) inspecting the predicted and observed profile shapes (read distributions) to ensure high profile similarity, and (3) inspecting DeepSHAP contribution scores on samples from held-out test chromosomes to ensure that predictions accurately highlight expected canonical motifs.

To estimate affinities for a sequence of interest, we then used an *in silico* marginalization approach (Affinity Distillation) in which we: (1) inserted a sequence of interest into at least 100 random backgrounds generated by sampling and randomly shuffling DNA sequences from held-out genomic peaks, (2) used the trained BPNet model to predict binding to these sequences, and (3) computed the mean marginal effect (marginalization score) of the insert on model output (here, sequencing read counts) relative to no insert (mean predicted log count ratio, or Δ log(counts)), thereby regressing to the mean of all context-dependent impacts and background effects (48–50).

To test if this *in silico* marginalization approach can accurately learn to predict quantitative thermodynamic affinities, we: (1) systematically trained BPNet on ChIP-seq data, (2) applied Affinity Distillation to predict affinities for sequences that had been characterized previously via *in vitro* thermodynamic assays, and then (3) quantified performance by calculating correlation coefficients and root mean squared errors (RMSE) between predicted and measured affinities. To test whether this neural network approach enhanced accuracy and/or precision relative to simpler models (51), we compared Affinity Distillation prediction performance with a variety of previously published best-in-class motif-based algorithms. We tested Affinity Distillation on iteratively more complex systems including systematic variations in nucleotides flanking binding sites of yeast TFs, systematic variations to core motifs, and general genomic context sequences in human TF systems.

### Affinity Distillation accurately extracts affinities from *in vitro* PB-exo yeast TF binding measurements

To assess whether AD can accurately learn quantitative binding affinities from chromatin immunoprecipitation (ChIP)-based approaches, we first tested AD on PB-exo data. PB-exo (34,52) generates genome-wide TF footprints by incubating a TF of interest with non-chromatinized DNA isolated from cells. After cross-linking, the DNA is digested with exonuclease, yielding high-resolution footprints of DNA sequence that is protected by TF binding. As PB-exo measurements take place *in vitro* where thermodynamic affinities are the sole determinant of TF occupancy, we expect Affinity Distillation to easily learn to accurately predict affinities. To quantitatively assess prediction accuracy, we turned to previously published BET-seq (25) data that used a microfluidic platform to quantify relative binding energies (ΔΔGs) for the *S. cerevisiae* model basic helix-loop-helix (bHLH) TFs Pho4 and Cbf1 interacting with all possible 5-nucleotide combinations flanking their known E-box consensus site (NNNNNCACGTGNNNNN, approximately 10^6^ double-stranded DNA (dsDNA) sequences) (**Fig. 1A**). As a baseline comparison between experiments, we first plotted log-transformed PB-exo read counts against BET-seq-measured ΔΔGs for the 884 and 294 NNNNNCACGTGNNNNN sequences that existed within the 100 bp windows of the PB-exo summits of Pho4 and Cbf1, respectively. Log-transformed PB-exo raw read counts were reasonably well-correlated with BET-seq-measured ΔΔGs over a 2-3 kcal/mol dynamic range (*r* =– 0.73, RMSE = 4.434 and *r* = –0.50, RMSE = 3.655 for Pho4 and Cbf1, respectively), as expected for a comparison between two somewhat similar *in vitro* experiments (**Fig. 1B**).

Next, we sought to test whether Affinity Distillation marginalization could accurately isolate energetic impacts of NNNNNCACGTGNNNNN sequences from PB-exo experiments and improve this correlation. We first trained the BPNet model to predict total counts and binding profiles for 4,558 of the 5,686 Pho4 binding regions and 846 of the 1,057 Cbf1 binding regions from the PB-exo data. Predicted binding profiles closely matched experimental observations, with contribution scores from DeepSHAP interpretations highlighting the expected E-box motifs underlying predicted areas of strong binding (**Fig. 1C**). Log-transformed count predictions correlated strongly with PB-exo experimental observations for both this training set (*r* = 0.91 and 0.81, respectively) and for remaining held-out regions from chromosomes 1 and 12 for Pho4 and chromosomes 5 and 6 for Cbf1 (*r* = 0.87 and 0.79, respectively) (**Fig. 1D**).

We then asked how Affinity Distillation marginalization scores correlated with BET-seq-measured ΔΔGs for the set of 884 Pho4 and 294 Cbf1 sequences present in both BET-seq and PB-exo datasets. The correlations were notably stronger than the baseline comparison (*r* = –0.91, RMSE = 1.772 and *r* = –0.74, RMSE = 2.063 for Pho4 and Cbf1, respectively). This suggests that *in silico* marginalization can denoise the counts associated with each match and account for various sources of experimental bias and variability to isolate thermodynamic impacts (**Fig. 1E**). To test whether the trained Affinity Distillation model can generalize beyond sequences in the PB-exo dataset to accurately predict affinities for as-yet-unseen sequences, we compared Affinity Distillation marginalization scores with ΔΔG measurements for all 1,048,576 possible NNNNNCACGTGNNNNN sequences in the BET-seq dataset, many of which are not found within the *S. cerevisiae* genome. Predictions remained tightly correlated with measurements (*r* =– 0.89 and –0.77 for Pho4 and Cbf1, respectively; **Fig. 1F**), establishing the generalizability of our approach. These results corroborate other reports that modern neural networks exhibit good generalization and a remarkably small difference between performance on training data and unforeseen examples (53,54), independent of the number of parameters in the network (55).

### Affinity Distillation predictions can be calibrated to yield absolute affinities

The raw predictions from the last layer of a neural network can be rich in information but their scale does not always have meaning (56). In this case, Affinity Distillation outputs marginalization scores that correlate with measured differences in binding energy but have no physical meaning on an absolute scale. Nevertheless, this observed linear correlation makes it possible to calibrate marginalization scores to binding free energies with only a small number of absolute affinity measurements (57,58). Using <1000 measurements sampled uniformly at random from the BET-seq dataset, we calibrated marginalization scores to derive ΔΔG values directly for all 1,048,576 sequences (*r* = 0.89, RMSE = 0.204 kcal/mol and *r* = 0.77, RMSE = 0.409 kcal/mol for Pho4 and Cbf1, respectively; see Methods; **Fig. 1G**). This calibration step can be generalized to any sequence-scoring method, allowing for direct comparison of prediction error (RMSE) while bypassing the need to compute motif similarity or alignment (59).

### Affinity Distillation predicts binding more accurately than other widely-used computational tools

To test if simpler algorithms could achieve the same performance, we predicted binding to the same 1,048,576 DNA sequences using three widely used best-in-class motif discovery tools that employ different modeling approaches: Weeder, an enumeration-based method shown to thoroughly cover the motif search space (60,61); the MEME suite member STREME, an optimization-based approach shown to perform well on ChIP-seq data (62); and MoDISco, a NN-based approach that uses importance scores to systematically summarize recurring predictive sequence patterns into consolidated motifs (46,63). Direct scoring is the best tool for assessment since it quantitatively reflects the similarity of *in vitro* binding preferences for TFs (59) and enables the computation of root mean squared errors in the same units as the dependent variable without relying on a particular matrix/motif similarity metric. In contrast to Affinity Distillation, which returns only a single high-confidence prediction and requires no parameter tuning beyond the original model training, these alternative approaches return multiple, often similar, motifs and it can be challenging to determine which returned motif is the most biophysically relevant (Supplement **Section 2**). To ensure a fair comparison, we scored all sequences in the library using the top 10 motifs returned by each algorithm (see Methods), calibrated scores using the same calibration process described above, and calculated an overall root mean squared error (RMSE) between predictions and measurements in absolute energetic units (kcal/mole).

Across all motifs, Affinity Distillation had a dramatically lower (*i.e.* more accurate) RMSE for Pho4 predictions (0.204 kcal/mol, corresponding to a 1.4-fold change in binding) than Weeder (mean RMSE = 0.396 for 10 motifs), STREME (mean RMSE = 0.433 for 8 motifs), and MoDISco (mean RMSE = 0.391 for 9 motifs) (**Fig. 1H**; Supplement **Section 3**). In addition, no single predicted Pho4 motif from any of these methods approached the accuracy of the Affinity Distillation model. Although improvements were more modest for Cbf1, Affinity Distillation still showed the greatest predictive accuracy (RMSE = 0.409 kcal/mol, compared with mean RMSE values of 0.507 from 9 motifs for Weeder, 0.581 from 7 motifs for STREME, and 0.610 from 9 motifs for MoDISco; **Fig. 1I**; Supplement **Section 3**). Together, these results establish that Affinity Distillation can accurately learn and generalize how changes to DNA sequence impact TF binding energies from *in vitro* data with greater accuracy than widely-used motif discovery methods and without the need to select which motif among the outputs is the most biophysically relevant.

### Affinity Distillation extracts affinities from models of *in-vivo* TF binding in yeast

In cells, extrinsic factors such as cooperativity and competition with other proteins and variations in local chromatin accessibility alter TF binding (64). If these extrinsic factors predominantly determine TF occupancy *in vivo*, we expect a significant drop in the ability of Affinity Distillation to successfully predict binding energies measured for a single TF interacting with DNA *in vitro*. To test this, we performed ChIP-nexus experiments with cross-linked TFs and DNA in yeast cells and then digested TF-bound DNA with an exonuclease to generate high-resolution binding footprints. As Pho4 is a TF that translocates to the nucleus and binds DNA in response to phosphate starvation, Pho4 experiments were performed under low phosphate conditions while Cbf1 experiments were performed under standard growth conditions. We obtained 1,621 and 1,174 genomic regions exhibiting statistically significant and reproducible enrichment of ChIP-nexus signals for Pho4 and Cbf1, respectively (Supplement **Fig. S8**; see Methods) (37).

We then trained BPNet neural network models on these *in vivo* data, predicted binding affinities via Affinity Distillation, and compared Affinity Distillation-predicted affinities with experimentally measured ΔΔGs (**Fig. 2A**). After training on 1,297 peaks and 939 peaks for Pho4 and Cbf1, BPNet-predicted log-transformed read counts for held-out chromosomes were well-correlated with experimental measurements (*r* = 0.66 and 0.72; **Fig. 2B**), predicted nucleotide-resolution binding profiles matched observed footprints, and DeepSHAP interpretations revealed canonical E-box motifs (**Fig. 2C**). As a baseline comparison between *in vivo* binding measurements and *in vitro* affinities, we again plotted raw log-transformed ChIP-nexus read counts against measured ΔΔGs for the 482 and 324 Pho4 and Cbf1 ChIP-nexus peaks containing particular NNNNNCACGTGNNNNN sequences measured within the BET-seq library. In comparison with PB-exo data, observed log-transformed ChIP-nexus raw counts were significantly more poorly correlated with BET-seq-measured relative free energies (r = –0.41, RMSE = 5.879 and r = –0.61, RMSE = 6.851 for Pho4 and Cbf1, respectively; **Figs. 2D, 1D**), particularly for Pho4, consistent with an increased importance of other extrinsic factors for properly localizing TFs in cells.

**Figure 2.**
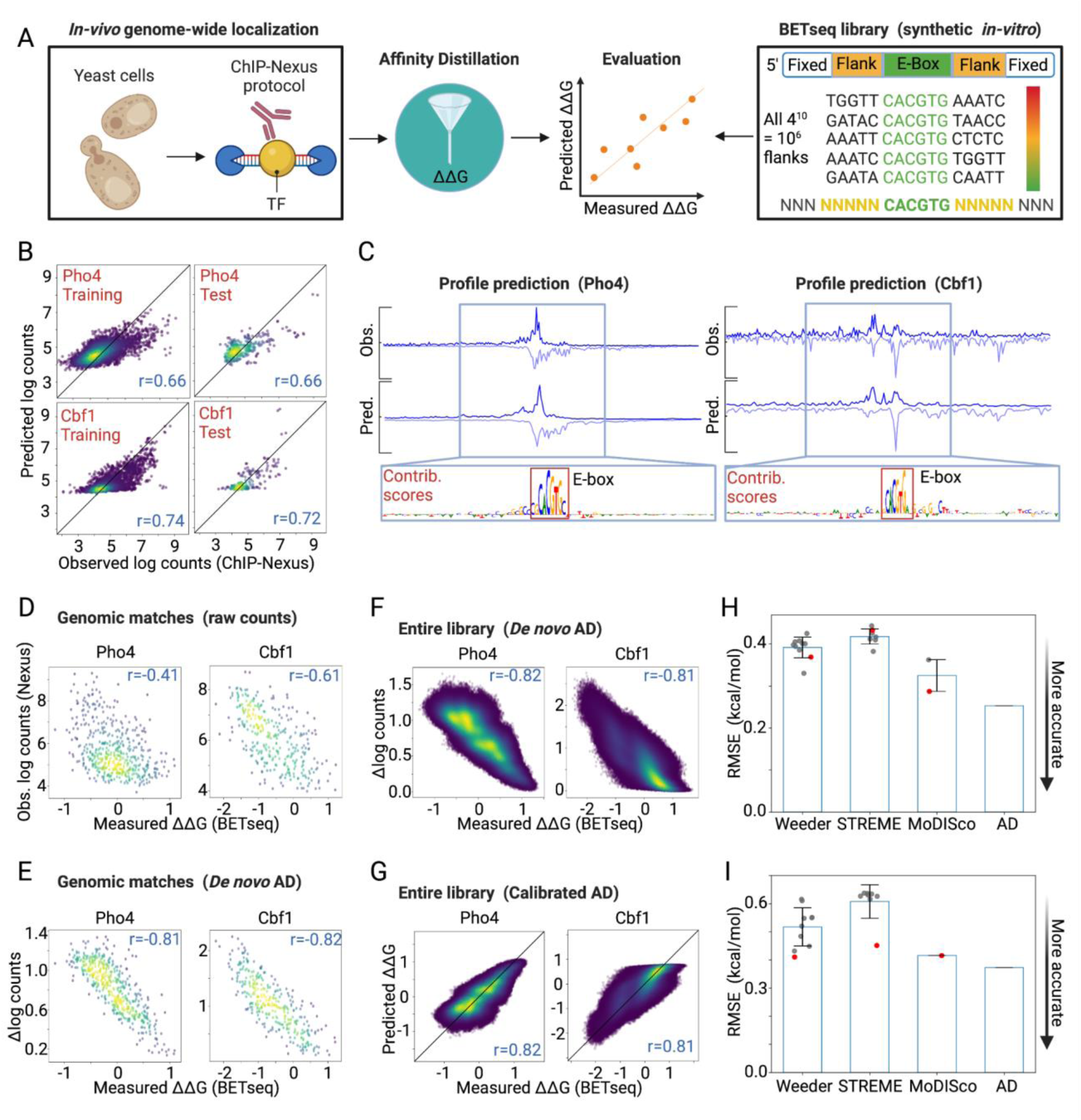
**Affinity Distillation extracts affinities from models of in-vivo TF binding in yeast.** (A) Pipeline overview: BPNet models are trained on ChIP-nexus data for Pho4 and Cbf1, Affinity Distillation (AD) extracts affinities, and AD outputs are compared with BET-seq-measured ΔΔGs. (B) Observed and predicted log-transformed read counts for Pho4 (top) and Cbf1 (bottom) for the training (left) and held-out test chromosomes (right). (C) Observed and predicted ChIP-nexus profiles for sample regions located on held-out test chromosomes; contribution scores highlight known CACGTG consensus sites within bound regions. (D) Measured log-transformed ChIP-nexus read counts vs. measured ΔΔGs for sequences present within both ChIP-nexus and BET-seq experiments (482 and 324 sequences for Pho4 and Cbf1). (E) AD-predicted marginalization scores vs. measured ΔΔGs for sequences present within both ChIP-nexus and BET-seq experiments. (F) AD-predicted marginalization scores vs. measured ΔΔGs for all 1,048,576 NNNNNCACGTGNNNNN sequences in BET-seq experiments. (G) Calibrated AD predictions vs measured ΔΔGs for all 1,048,576 NNNNNCACGTGNNNNN sequences in BET-seq experiments. (H) Post-calibration RMSEs of predictions vs observations for Pho4. Gray markers indicate individual motif performances; red markers indicate the performance of the top output of each algorithm; blue bars indicate mean values; error bars indicate standard deviation. (I) Same as H for Cbf1.

Despite a significant drop in the correlation between raw ChIP-nexus read counts and measured affinities, Affinity Distillation remained capable of accurately predicting binding affinities for both genomic matches present in both datasets (r = –0.81, RMSE = 1.092 and r = –0.82, RMSE = 1.970 for Pho4 and Cbf1, respectively) (**Fig. 2E**) or for all 1,048,576 possible NNNNNCACGTGNNNNN sequences (r = 0.82, RMSE = 0.253 and r = 0.81, RMSE = 0.373 for Pho4 and Cbf1, respectively) (**Fig. 2F**), with this strong linear correlation facilitating calibration of predicted marginalization scores into energy space.

Affinity Distillation again predicted *in vitro* affinities more accurately than Weeder (mean RMSE = 0.392 and 0.517 for Pho4 and Cbf1), STREME (mean RMSE = 0.418 and 0.608 for Pho4 and Cbf1), and MoDISco (mean RMSE = 0.325 and 0.416 for Pho4 and Cbf1) (**Figs. 2H,I**; Supplement **Section 3**). Together, these results establish that Affinity Distillation was able to successfully extract affinities from TF binding *in vivo* even when the observed raw *in vivo* signal for the native genomic regions was only moderately correlated with measured affinities.

### Affinity Distillation can decipher the differential specificity of paralogous TFs

As a next test of Affinity Distillation’s performance, we sought to determine whether the algorithm could correctly predict differential binding for paralogous TFs that bind very similar motifs yet target distinct genes *in vivo* (**Fig. 3A**). Although Pho4 and Cbf1 share a similar E-box motif (**Fig. 3B**), they perform different functions in cells: phosphate starvation leads to dephosphorylation and nuclear localization of Pho4, which drives the expression of genes associated with phosphate scavenging; Cbf1 is upregulated under replication stress and associates with other TFs to activate or repress a wide variety of downstream target genes (29,65–68). Consistent with these distinct physiological roles, only a small fraction of peaks were bound by both TFs in the ChIP-nexus data, where only Pho4 was profiled under phosphate starved conditions (**Fig. 3C**). In prior ChIP-seq experiments, where both Cbf1 and Pho4 were tested under phosphate starved conditions, more than half of the accessible consensus ‘CACGTG’ sites were preferentially bound by Cbf1 alone or by both proteins (**Fig. 3D**) (68). This unique localization is thought to result, at least in part, from different intrinsic affinities to genomic regions (29,30) such that Cbf1 can outcompete Pho4 at consensus genomic loci that would otherwise be Pho4-occupied (29,69). Here, we investigated whether Affinity Distillation can learn to predict subtle binding affinity and specificity differences between TF paralogs by training the model on Pho4 and Cbf1 *in vivo* ChIP-nexus data and then attempting to predict differential Pho4 and Cbf1 binding observed in *in vitro* genomic context protein binding microarray (gcPBM) experiments. gcPBM experiments quantify binding of fluorescently-labeled Pho4 or Cbf1 proteins to 20,414 surface-immobilized 60 nucleotide dsDNA probes, each of which contains a central 36-nucleotide sequence corresponding to a genomic locus bound by either Pho4, Cbf1, or both in cells; prior direct comparisons between measured affinities and gcPBM probe intensities have established that log-transformed probe intensities correlate with absolute binding affinities over a wide dynamic range (32,70,71).

**Figure 3.**
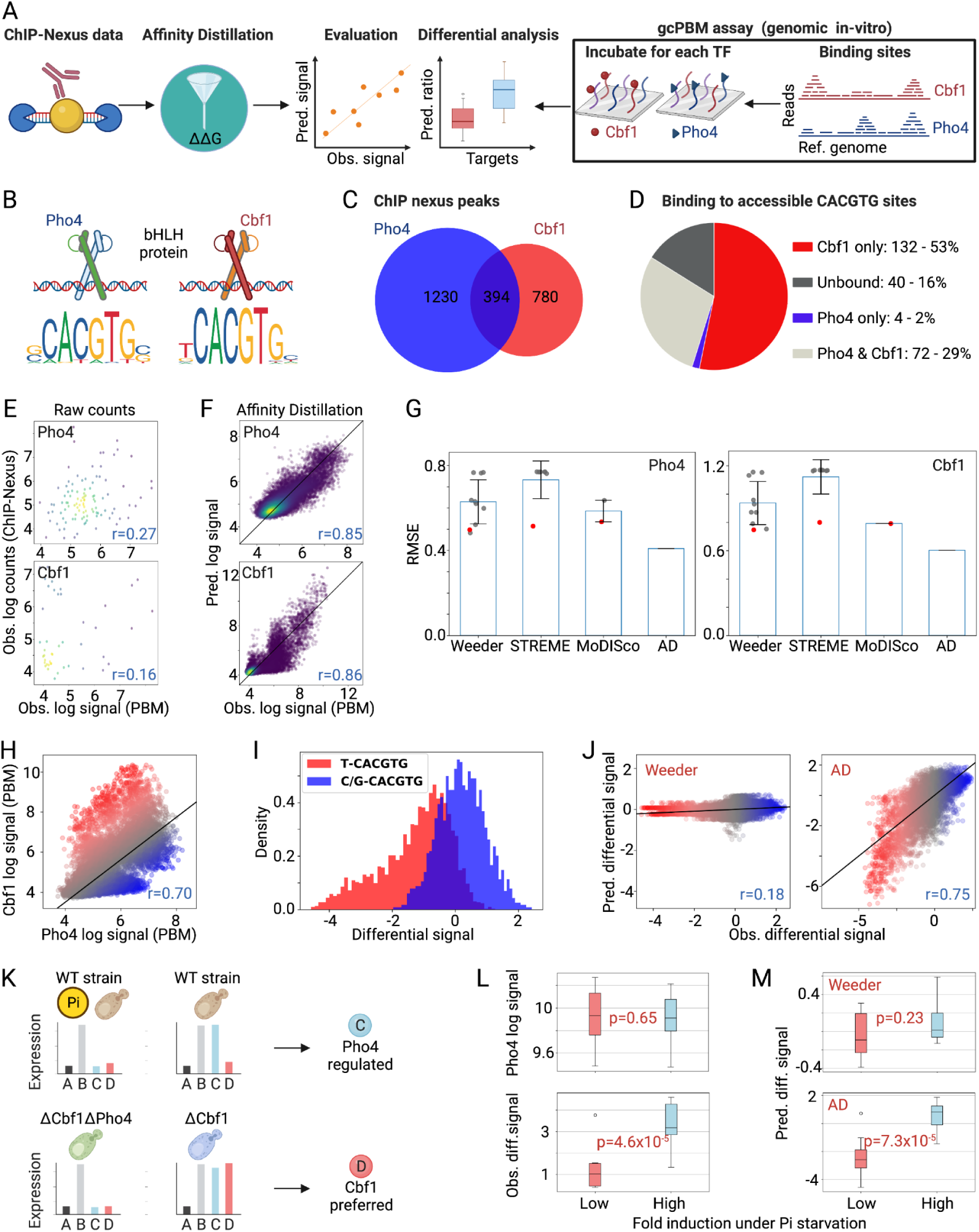
**Affinity Distillation can decipher the differential specificity of paralogous TFs.** (A) Pipeline overview: BPNet models are trained on ChIP-nexus data for Pho4 and Cbf1, Affinity Distillation (AD) extracts affinities, and AD outputs are compared with gcPBM-measured intensities (29). (B) Cbf1 and Pho4 position weight matrix (PWM) models from JASPAR (129). (C) Venn diagram of ChIP-nexus peaks of Cbf1 and Pho4 (see Methods). (D) Pie chart showing Pho4 and Cbf1 binding at all accessible consensus binding sites from ChIP-seq data in no-Pi conditions as previously reported (68). (E) Observed log-transformed ChIP-nexus counts vs. observed log-transformed gcPBM intensities for genomic loci bound in both ChIP-nexus and gcPBM experiments (104 and 44 sequences for Pho4 and Cbf1). (F) Calibrated AD-predicted log intensities vs. observed log-transformed intensities for all sequences within gcPBM libraries (20,414 sequences). (G) Post-calibration RMSEs of predictions vs observations. Gray markers indicate individual motif performances; red markers indicate the performance of the top output of each algorithm; blue bars indicate mean values; error bars indicate standard deviation. (H) Measured gcPBM probe intensities for Cbf1 vs. Pho4 (29); red and blue markers indicate Cbf1-and Pho4-preferred sequences, quantified by residuals from the regression line. (I) Histogram of differential signals for sequences with a 5’ T nucleotide (red, consistent with Cbf1 preferences) or a 5’ C or G (blue, consistent with Pho4 preferences) preceding a CACGTG motif. (J) Predicted vs. observed differential signal for Weeder (left) and AD (right); line indicates linear regression. (K) Schematic showing behavior of Pho4-regulated (blue) genes and Cbf1-preferred (red) genes as classified in (29,68) under different conditions; ΔCbf1 and ΔCbf1ΔPho4 signify deletion strains. (L) Box plots showing observed in vitro Pho4 log signal intensities (top) and Pho4/Cbf1 differential intensities (bottom) for genes with low versus high fold induction in response to Pi starvation. (M) Box plots showing predicted differential signals from Weeder2 (top) and AD (bottom) for the same genes.

Observed ChIP-nexus log-transformed counts for Pho4 and Cbf1 were relatively poorly correlated with observed log-transformed gcPBM intensities (*r* = 0.27 and *r* = 0.16 for Pho4 and Cbf1; **Fig. 3E**). Nevertheless, Affinity Distillation predictions after training on the same data again showed dramatically improved accuracy (*r* = 0.85, RMSE = 0.409 and *r* = 0.86, RMSE = 0.604 for Pho4 and Cbf1; **Fig. 3F**). As with previous test applications (**Fig. 2**), the RMSE between predictions and observations was significantly lower (*i.e.* more accurate) for Affinity Distillation than for the Weeder2, STREME and MoDISco motif-prediction algorithms without a need to select from a range of potential motifs (**Fig. 3G**; Supplement **Section 3**).

Next, we tested whether Affinity Distillation could accurately predict *differential* binding preferences between these paralogs. Comparing measured per-probe intensities from Pho4 and Cbf1 gcPBM experiments reveals three distinct populations: probes equally bound by both proteins (**Fig. 3H**, gray), probes preferentially bound by Cbf1 (**Fig. 3H**, red), and probes preferentially bound by Pho4 (**Fig. 3H**, blue). Here, we quantify a paralog-specific ‘differential signal’ as the residual from the best fit line for Cbf1 vs. Pho4 log-transformed intensities such that positive and negative differential signals report on the degree to which a given probe is preferentially bound by either Pho4 or Cbf1, respectively.

Consistent with prior work (22,68), sequences with a T nucleotide 5’ to the CACGTG consensus are Cbf1-preferred (lower differential signal) and sequences with a C/G nucleotide 5’ to the CACGTG consensus are Pho4-preferred (higher differential signal), with the mean differential signals significantly different between the two groups (p < 1×10^-6^ evaluated using Welch’s T-test for the means of two independent samples of scores with unequal variances; **Fig. 3I**). Although Weeder2 had been the best-performing alternative algorithm in predicting binding of either protein in isolation (**Fig. 3G**), it was unable to accurately distinguish probes preferred by one paralog over another (*r* = 0.18; **Fig. 3J**). By contrast, Affinity Distillation accurately identified paralog-preferred genomic loci (*r* = 0.75; **Fig. 3J**).

### Affinity Distillation can improve the differential classification of regulatory targets

To test if an enhanced ability to predict paralog-preferred loci improves the identification of regulatory networks in cells, we turned to previously published Pho4 gene expression data from wildtype cells (WT) and cells lacking Cbf1 (ΔCbf1) collected under high and no phosphate conditions (as genes regulated by Pho4 binding are expressed only upon phosphate starvation) (68). These data revealed 4 classes of genes: (1) Pho4-regulated genes (**Fig. 3K**, blue), which were activated by Pho4 only under no phosphate conditions in the wildtype cells; (2) Cbf1-preferred genes (**Fig. 3K**, red), which were activated by Pho4 only when Cbf1 is absent (in ΔCbf1 but not WT); (3) always active genes (**Fig. 3K**, light gray), which were expressed in all cases; and (4) inactive genes (**Fig. 3K**, black), which were never expressed (see Methods) (29). If Affinity Distillation’s ability to discriminate Pho4-regulated from Cbf1-preferred binding sites allows more accurate recapitulation of *in vivo* regulatory networks, we expect Affinity Distillation-predicted differential signals to accurately identify *bona fide* Pho4-regulated targets.

Even though the observed gcPBM Pho4 signal was not significantly different between genomic loci classified as Pho4-regulated or Cbf1-preferred (**Fig. 3L**), the observed differential signal was significantly higher for Pho4-regulated genes (evaluated using Welch’s T-test for the means of two independent samples of scores with unequal variances; **Fig. 3L**). Consistent with previous observations that Weeder cannot accurately discriminate Pho4-from Cbf1-preferred loci (**Fig. 3J**), Weeder-predicted intensities did not differ between the two classes (**Fig. 3M**). Nevertheless, Affinity Distillation-predicted differential intensities clearly distinguished the two classes (p = 7.3×10^-5^;**Fig. 3M**), establishing that an enhanced ability to learn context-dependent preferences can provide physiologically-relevant insights into regulatory networks.

### Affinity Distillation enhances prediction accuracy by matching the dynamic range of measured affinities

We hypothesized that Affinity Distillation likely outperforms traditional motif-based representation models because it captures a high-dimensional representation of the binding landscape that accounts for the fact that TFs recognize distinct binding sites with a wide range of affinities via different binding mechanisms and protein conformations (24,32,72–74), many of which are not accurately modeled by a motif or PWM. To test this, we plotted the distribution of log-transformed gcPBM intensities for Pho4 (**Fig. 4B**, blue) along with predicted log-transformed intensities from Affinity Distillation, STREME, MoDisco, and Weeder trained on Pho4 ChIP-nexus data (**Fig. 4B**, orange). The experimentally observed log-transformed gcPBM intensity distribution is composed of a large approximately Lorentzian low-intensity peak (representing low-affinity background binding) and a high-intensity tail of specific binding spanning 3 orders of magnitude (**Fig. 4B**, blue). Of the 4 predicted distributions, Affinity Distillation’s distribution most closely matched this observed distribution, with a single background binding peak and a high-affinity tail extending over the largest dynamic range (log-transformed standard deviation of 0.77 for the observed distribution *vs.* 0.66, 0.63, 0.61, and 0.60 for AD, Weeder, MoDISco, and STREME predictions, respectively) (**Fig. 4B,C**). By contrast, predictions for the motif-based predictions were truncated at high affinities, regardless of the size of the PWM. Further suggesting that a wide dynamic range is essential for accurate binding predictions, the overall root mean squared error of predictions was negatively correlated with the observed dynamic range (**Fig. 4C**). *In silico* marginalization against different sequence backgrounds appeared to aid in maximizing dynamic range, as dynamic range increased with the number of backgrounds (**Fig. 4D**).

**Figure 4.**
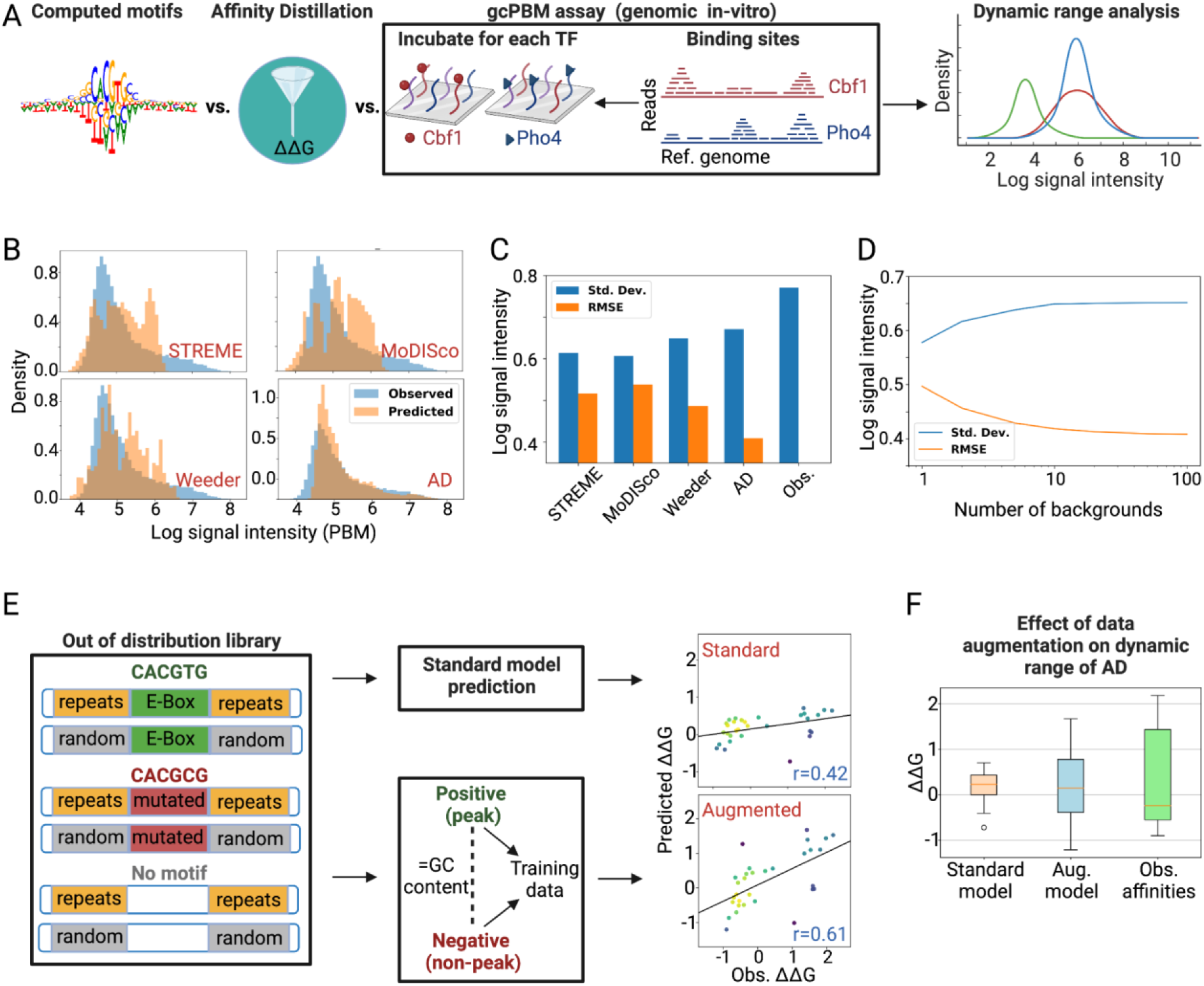
**Affinity Distillation predictions recapitulate the dynamic range of observed affinities.** (A) Pipeline overview: Motif-based methods and Affinity Distillation (AD) are trained on Pho4 in vivo ChIP nexus data and used to predict affinities for sequences in *in vitro* gcPBM experiments; predicted affinities are then compared with experimental observations. (B) Histograms showing observed (blue) and predicted (orange) intensity distributions for STREME, MoDISco, Weeder and AD. (C) Observed and predicted standard deviations and RMSEs for log-transformed intensity distributions. (D) AD-predictions’ standard deviations and RMSEs as a function of the number of backgrounds (1, 2, 5, 10, 20, 50, 100 seqs) used in in silico marginalization. (E) Schematic of GC-matched augmentation strategy testing prediction accuracy for an out-of-distribution yeast DNA library designed to test how short tandem repeats flanking known binding sites alter binding affinities (32); comparisons between AD predictions and observed ΔΔGs for standard and augmented models are shown at right. (F) Box plots showing dynamic range of observed and AD-predicted ΔΔGs.

Responsible development and dissemination of powerful algorithms requires transparently evaluating prediction accuracy under conditions where the algorithm is likely to perform well and where performance may break down. To test the degree to which Affinity Distillation can learn binding mechanisms and contexts that do not exist in the yeast genome, we attempted to use the trained model to predict Pho4 preferences to all possible short tandem repeats (STRs), a recently-described novel class of cis-regulatory element (31,32,75). While the same model architecture successfully learned to predict impacts of STRs on binding for mammalian bHLH TFs, STRs are shorter and less frequent in the compact *S. cerevisiae* genome, making the task significantly more challenging. As expected for out-of-distribution effects, applying the standard training protocol to a Pho4 ChIP-nexus model led to poor predictive performance (r=0.42; **Fig. 4E**).

To improve performance, we turned to data augmentation, which can mitigate a large search space (76) and even synthetically generate realistic biological signals (77), by adding negative examples to the dataset. For each peak in the training set, we randomly selected a negative region (at least 1 kb outside genomic peaks) with the same GC content (within 5%). To maintain a 10:1 ratio of positives to negatives in the training set, we then jittered the positive region (peak) 10 times up to 200 bp away from the summit. This data augmentation strategy substantially expanded the dynamic range of the predictions and increased prediction accuracy (r=0.61; **Fig. 4E,F**), suggesting a generalizable approach for improving performance when attempting to predict affinities for out-of-distribution sequences.

### Affinity Distillation extracts affinities from mammalian ChIP-seq models

The yeast genome is relatively small, with gene expression controlled primarily by TF binding to gene-proximal promoters (78), and thermodynamic models based on equilibrium binding affinities have previously explained the majority of observed variance in gene expression (79). Therefore, it is perhaps not surprising that Affinity Distillation can accurately extract quantitative binding affinities from *in vivo* Tf binding data in yeast. Regulation of transcription in mammalian cells is thought to be significantly more complex, depending on transient interactions between promoters and distal enhancers that are in turn regulated by the formation of 3D chromatin compartments (80). Here, we tested whether Affinity Distillation could still recover quantitative binding affinities for the human glucocorticoid receptor (GR) even as the relative importance of extrinsic factors in correct localization increases (**Fig. 5A**).

**Figure 5.**
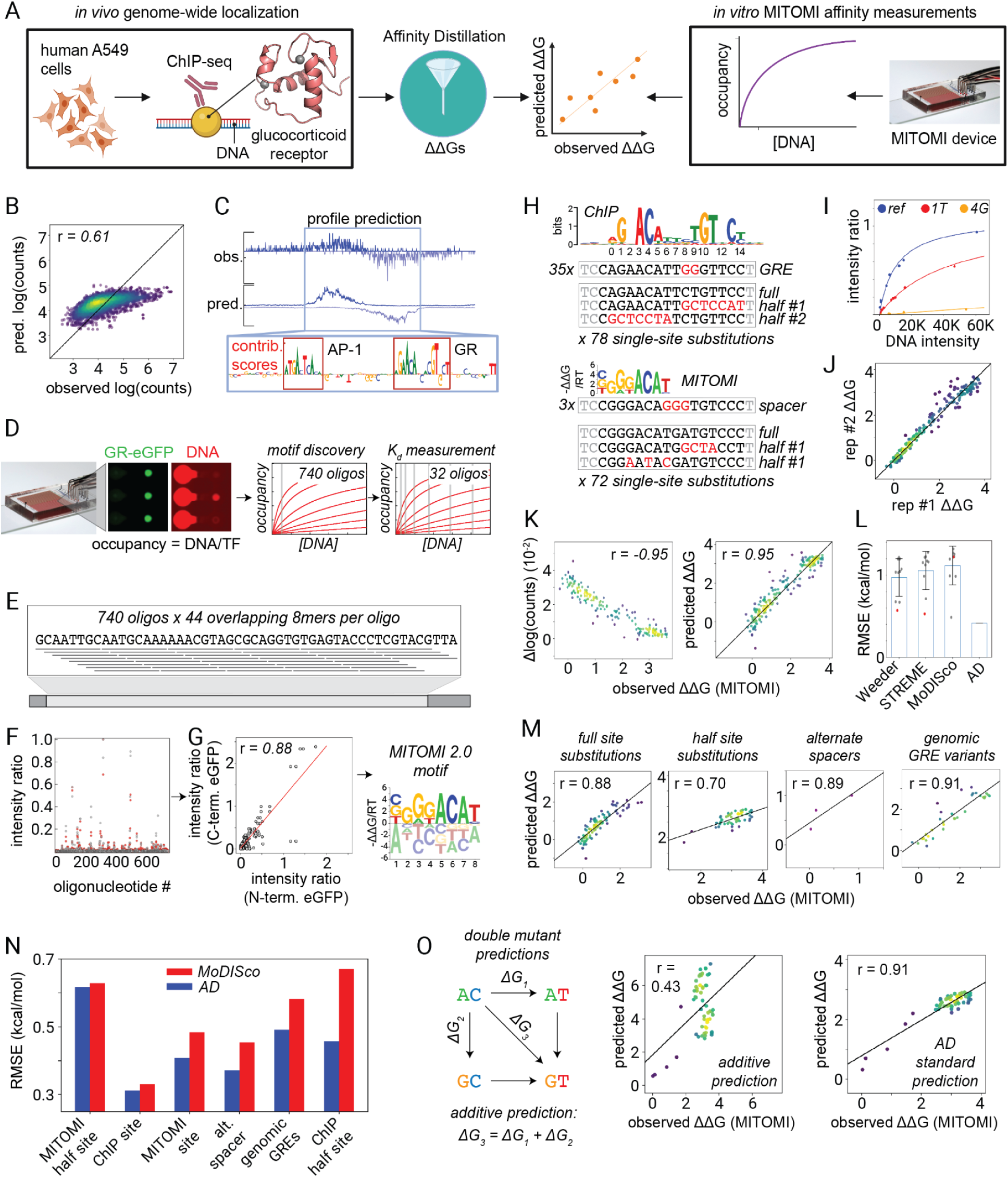
**Affinity Distillation can extract human GR binding affinities from ChIPseq data.** (A) Pipeline overview: BPNet is trained on ChIP-seq data for GR [pdb: 1gdc], Affinity Distillation (AD) extracts affinities, and AD outputs are compared with MITOMI-measured ΔΔGs. (B) Observed and predicted log-transformed read counts for GR on held-out test data. (C) Observed and predicted ChIP-seq profiles for sample regions located on held-out test chromosomes; contribution scores highlight GR and AP-1 consensus sites within bound regions. (D) Overview of MITOMI experiments: first, relative fluorescence intensities measured for pseudorandom sequences at a single concentration reveal preferred half sites (MITOMI 2.0); subsequent concentration-dependent binding measurements of systematically varying DNA libraries quantify Kds (MITOMI). (E) Schematic of pseudorandom 8-mer library quantified by MITOMI 2.0 comprised of 740 70 bp dsDNA sequences with 44 overlapping 8-mers per sequenceA. (F) Scatterplot showing measured intensity ratios (DNA/transcription factor) for each dsDNA sequence in the pseudorandom 8-mer library. (G) Scatterplot showing normalized binding intensity ratios for GR constructs with either C-or N-terminal eGFP tags interacting with the pseudorandom 8-mer library and motif derived from these ratios. (H) Overview of MITOMI library including dimeric versions of the optimized MITOMI-derived monomer, the ChIP-derived site, and systematic variations to half and full sites (188 sequences total). (I) Representative concentration-dependent binding for GR binding to DNA sequences containing an intact motif (Reference (Ref.); GGGACATGATGTCCC), a mutated flanking sequence (1T; GTGACATGATGTCCC), or a mutated motif (4G; GGGAGATGATGTCCC) (see Supplement for all other binding curves). (J) Scatterplot showing replicate MITOMI ΔΔG measurements. (K) Scatterplots showing de novo marginalization scores (left) and post-calibration predicted affinities (right) vs. measured ΔΔGs. (L) Post-calibration root mean squared error (RMSE) of predictions vs observations. Gray markers indicate individual motif performances; red markers indicate the performance of the top output of each algorithm; blue bars indicate mean values; error bars indicate standard deviation. (M) Scatterplots showing the predictive performance of Affinity Distillation on single substitution variations of consensus sites (MITOMI and ChIP), mutations to half sites (MITOMI and ChIP), alternate spacer sequences, and genomic GRE variants. (N) Breakdown of RMSE by type of variation in the library, sorted by difference between Affinity Distillation and the best performing motif (in this case MoDISco). (O) Schematic of a double mutant cycle (left) and scatterplots showing the performance of a simulated additive AD prediction and standard AD prediction (right) for MITOMI probes containing multiple mutations from the consensus site.

Glucocorticoid receptor (GR) is a broadly-expressed nuclear hormone receptor (NHR) TF containing a DNA binding domain, a ligand binding domain, and an activation domain. Upon binding glucocorticoid hormones, GR translocates to the nucleus and upregulates a wide variety of developmental, metabolic, and immune-associated transcriptional programs (81,82). Using the irreproducible discovery rate (IDR) framework (83) to identify reproducible peaks within a prior ENCODE ChIP-seq dataset yielded 17,203 GR-bound peaks, many of which contained canonical glucocorticoid binding sites (GBSs) comprised of inverted GNACA repeats typically separated by a 3 nt degenerate spacer. As ChIP-seq GR experiments returned many bound peaks and we anticipate an increased influence of extrinsic factors on binding, we trained a larger capacity model (see Methods) on 13,677 peaks after withholding regions from chromosomes 8 and 9 (10% of total peaks) as a test set and regions from chromosomes 16, 17, and 18 (10% of total peaks) as a validation set.

The model predicted *in vivo* GR binding with a slightly reduced accuracy as compared to *S. cerevisiae* Pho4 and Cbf1 (**Fig. 5B**; *r* = 0.61 for GR held-out test data, compared with *r* = 0.66 and 0.72 for Pho4 and Cbf1 tests) and contribution scores for ChIP peaks often revealed motifs associated with other TFs, including the AP-1 motif (84), highlighting that cooperation with other TFs is essential for proper GR localization in cells (**Fig. 5C**).

In parallel, we used the MITOMI microfluidic platform (22) to quantify affinities (**Fig. 5D**) for recombinantly expressed and purified C-terminally eGFP-tagged GR interacting with two different DNA libraries. To enable unbiased identification of GR binding specificities, we first profiled GR binding at a single low concentration to a de Bruijn dsDNA library of 740 pseudorandom sequences containing all possible 8-mer DNA combinations arranged in a minimally compact space (85) (**Fig. 5E**), which returns intensity information linearly correlated with relative differences in binding energy (86). Although this pseudorandom 8-mer library did not contain any true GNACANNNTGTNC dimeric sites, these measurements revealed a strongly preferred monomeric half site (**Fig. 5F,G**), consistent with prior reports that both monomeric and dimeric forms of GR bind *in vivo* to regulate distinct expression programs (87,88). The MITOMI-derived motif closely resembled the ChIP-derived GBS (89) but with a preference for GGG instead of AGA upstream of the canonical ACA (see Supplement).

We then profiled concentration-dependent GR binding to quantify absolute affinities (*K*_d_s) for an additional 188 dsDNA sequences containing systematic single-nucleotide substitutions within monomeric and dimeric versions of the MITOMI-and ChIP-derived high affinity sites (**Fig. 5H**), synthetic GBSs with variable half-site spacings, and several naturally-occurring glucocorticoid response elements (GREs) (see Supplement GR tables). Concentration-dependent binding measurements for each sequence were globally fit to a Langmuir isotherm to extract *K*_d_s (assuming binding stoichiometry remained unchanged across sequences); all ΔΔGs were calculated relative to the ChIP consensus site (**Fig. 5I**). Calculated ΔΔGs were again highly reproducible across replicates (*r* = 0.98; **Fig. 5J**). Consistent with MITOMI pseudorandom library results and with prior observations that many TF sites are optimized for moderate affinities *in vivo* to facilitate dynamic regulation (90–97), mutations to the ChIP-derived consensus that yielded a more symmetric and MITOMI-like motif consistently increased affinities (See Supplemental GR table).

As in previous comparisons, Affinity Distillation accurately predicted ΔΔGs for the entire MITOMI library on par with replicates (r = 0.95; **Fig. 5K**), outperforming Weeder, STREME, and MoDISco, with an RMSE of 0.414 kcal/mol compared to mean RMSEs of 0.971 kcal/mol, 1.06 kcal/mol, and 1.12 kcal/mol, respectively (**Fig. 5L**).

Across all 4 library types (single site substitutions in the dimeric and monomeric sites, alternate half site spacers, and GR-bound genomic loci), Affinity Distillation outperformed the best-performing (lowest RMSE) alternative algorithm (MoDISco), with the largest performance boost for ChIP-half sites and naturally-occurring genomic glucocorticoid response elements (GREs) that had multiple mutations from the consensus site (**Fig. 5M, N**). To explicitly test whether Affinity Distillation’s increased accuracy derives from an ability to learn epistatic impacts of nonadditive mutations, we compared predicted Affinity Distillation additive predictions and Affinity Distillation-predicted ΔΔGs with measured values for libraries of sequences in which pairs of positions were mutated alone and in combination (**Fig. 5O**; left). While additive models continued to predict increasingly deleterious impacts even beyond levels of non-specific binding (**Fig. 5O**; middle), Affinity Distillation successfully recapitulated the non-additive effects of multiple binding site mutations (**Fig. 5O**; right).

### Affinity Distillation performance is generalizable across multiple mammalian TF classes and cell types

Thus far, we have established that Affinity Distillation can extract thermodynamic binding information from Pho4, Cbf1, and GR *in vivo* genome-wide occupancies. However, as the GR NHR binds a relatively long binding site with high affinity, thermodynamic binding affinities are more likely to play a dominant role in dictating localization. To test whether Affinity Distillation can accurately recover affinities across a wide variety of additional TF structural classes and human cell types, we turned to publicly available ENCODE ChIP-seq data to train additional TF-specific models for TFs across the bHLH, ETS, and E2F TF families: MAX in HeLa-S3 cells (bHLH; 20438 peak regions), c-Myc in HeLa-S3 and K562 cells (bHLH; 16391 and 17318 peak regions), GABPA in HepG2 cells and liver tissue (ETS; 6129 and 9543 peak regions), and E2F1 in MCF7 cells (E2F; 19373 peak regions).

We then attempted to predict affinities for up to 30,000 gcPBM probes for each TF or TF family (30) and assessed predictive performance (**Fig. 6A**). Across all 6 TFs, ChIP-seq counts were only weakly correlated with measured signal intensities for the same genomic sequences in gcPBM experiments, with Pearson correlation coefficients ranging from 0.18 (MAX and myc) to 0.31 (E2F1 with a very limited number of probes).

**Figure 6.**
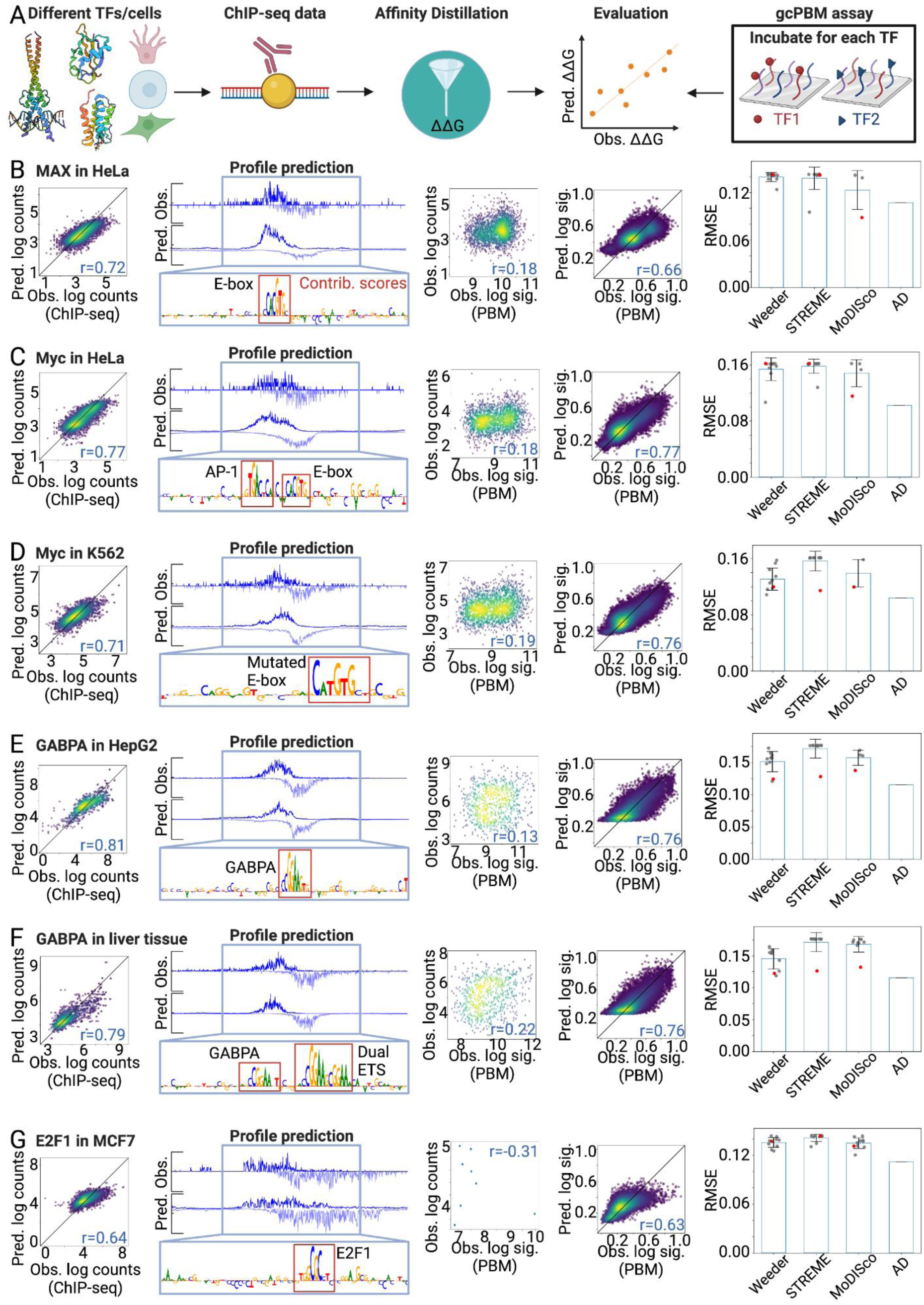
**Affinity Distillation can extract affinities of mammalian genomic binding sites.** (A) Pipeline overview: BPNet models are trained on ChIP-seq data of different TF structural families [pdb: 5EYO, 2JUO, 6G0P] in different cell types, Affinity Distillation (AD) extracts the affinities, and AD outputs are compared with gcPBM intensities. (B) Panel 1: Observed and predicted read coverage across peaks of held-out chromosomes for MAX model in HeLa-S3 cells; Panel 2: observed and predicted ChIP-seq profiles for a sample region located on held-out test chromosomes; Panel 3: log-transformed ChIPseq read counts and gcPBM intensities for sequences present within both ChIP-seq and gcPBM experiments; Panel 4: post-calibration AD-predicted affinities vs. gcPBM intensities (30); Panel 5: post-calibration root mean squared errors (RMSE) for predictions vs observations. Gray markers indicate individual motif performances; red markers indicate the performance of the top output of each algorithm; blue bars indicate mean values; error bars indicate standard deviation. (C–G) Same as B for Myc in HeLa-S3, Myc in K562, GABPA in HepG2, GABPA in liver tissue, and E2F1 in MCF7, respectively.

As a first test of whether Affinity Distillation could more generally learn thermodynamic affinities from *in vivo* ChIP-seq data, we analyzed data for MAX, a bHLH TF that binds to the palindromic CACGTG E-box motif either as a homodimer or as a heterodimer with other bHLH proteins such as Myc (98) (**Fig. 6B**). MAX-MAX homodimers are transcriptionally inert, while Myc-MAX heterodimers drive strong transcriptional activation by promoting elongation (99–101); as a result, dysregulation of Myc can drive oncogenic progression in many human cancers, most commonly through Myc overexpression (102). After training on 16,324 peaks from MAX ChIP-seq, predicted log-transformed read counts for held-out chromosomes were well-correlated with experimental measurements (r = 0.72). The predicted binding profiles matched observed footprints in cells, and DeepSHAP interpretations revealed canonical E-box motifs within bound regions (**Fig. 6B**). Post training, Affinity Distillation was capable of accurately predicting binding affinities for the entire gcPBM library and again outperformed all other techniques (r = 0.66; **Fig. 6B**, and **Fig. S9**).

Models trained on 12833 peaks from Myc ChIP-seq data were similarly successful in predicting held-out read counts and observed binding profiles for two very different cell types (r = 0.77 and 0.71 for adherent HeLa cervical cancer cells and suspension K562 lymphoblastoid cells, respectively) (**Fig. 6C,D**). Affinity Distillation was capable of accurately predicting binding affinities for the entire gcPBM library after training on two very different cell types (r = 0.77 and 0.76), again outperforming all motif-based methods tested (**Figs. 6C,D**).

Next, we investigated GABPA, an ETS family TF that recognizes GGA(A/T) consensus sequences via a winged helix-turn-helix structure. In addition to being required for GABPB recruitment, GABPA is also thought to be a key transcriptional regulator in cell cycle control and metabolism in myeloid cells and other cell types (103) (**Fig. 6E**). As for the other TFs, Affinity Distillation was able to accurately predict binding affinities for the entire gcPBM library (r = 0.76 and 0.76). A final comparison for E2F1, a member of the E2F family with well-documented roles in cell cycle progression and apoptosis, as well as newly-emerging roles in metabolic homeostasis and obesity (104), again predicted probe intensities with high accuracy despite there being only 8 shared sequences and poor agreement between ChIP-seq derived counts and gcPBM intensities (see Methods) (**Fig. 6F**).

## Discussion

Traditionally, extracting affinities from *in vivo* assays has focused on discovering short motifs enriched within bound loci and then using these motif-based models to predict DNA binding with somewhat limited success (105–107). By contrast, modern machine learning methods such as deep neural networks can predict binding with high accuracy but are sometimes dismissed as overparameterized black box models with no way to extract biophysical information (108). To remedy this, some studies have suggested building stereotyped networks with fixed architectures, sacrificing flexibility in modeling and training to obtain nodes and weights that have explicit biophysical interpretations (109,110). Here, we establish that with appropriate correction of experimental biases (Supplement **Section 6**), deep neural networks trained to predict TF binding profiles from ChIP data can learn to predict thermodynamic affinities for a wide variety of genomic sequences without *a priori* restricting architectures or weights.

The development of neural networks capable of accurately predicting binding affinities for novel sequences opens up opportunities to perform *in silico* biophysics experiments at unprecedented scale. There are >10^60^ possible nucleotide combinations for a 100 bp cis-regulatory element, dwarfing the sequence space that can be probed by even the highest throughput experiments. The neural network model presented here can predict relative binding affinities for millions of sequences per day; moreover, these predictions can be calibrated to actual energies with fewer than one thousand *in vitro* measurements. Affinity Distillation can therefore serve as an *in silico* biophysical oracle capable of predicting how systematic variations to central target sites or local sequence context impact binding. Indeed, Affinity Distillation has already been used to validate the role of short tandem repeats in TF recruitment (32). Though used for validation, the short tandem repeat phenomenon could have in theory been discovered through neural networks. In future work, Affinity Distillation could identify cooperative regulatory modules by testing the impacts of mutating individual motifs alone and in tandem. All of these analyses have been possible with base-resolution networks, but with Affinity Distillation, they can be done in the language of thermodynamics.

Affinity Distillation will be particularly useful in predicting binding to low-affinity DNA-binding sites that can be critical for gene regulation (33). As a specific example, developmental enhancers with low-affinity binding sites can mediate robust patterns of gene expression when they are organized with optimal syntax (111). Accurately predicting binding in these cases is particularly challenging using current techniques, as non-canonical binding motifs are typically disregarded by motif-based methods. As another example, Affinity Distillation can predict differential regulation by proteins with extremely similar DNA-binding domains that share high-affinity binding preferences but have distinct DNA-binding profiles and preferences for low affinity sites (112). Such paralog-specific preferences cause difficulties for motif-based models, as the models frequently characterize the common binding sites but fail to capture the sites specific to each TF. We have shown that Affinity Distillation is capable of learning paralog-specific affinities, enabling us to distinguish between true physiological targets from targets that are blocked off by competitive paralogs. Overall, Affinity Distillation matches the paralog-specific preferences and the overall dynamic range of observed affinities better than all methods that we investigated.

## Conclusion

In this work, we demonstrated Affinity Distillation, a method that bridges the gap between modern neural network models capable of predicting *in vivo* binding without any mechanistic understanding and ground-up biophysical thermodynamic models of TF-DNA interactions. Using an *in silico* marginalization approach developed to interrogate deep learning models, Affinity Distillation learns accurate affinities of binding sites, flank variations, genomic sequences, and even paralog-specific affinities, enabling in silico differential analyses between related TFs. By providing biophysical interpretations of deep learning models, Affinity Distillation makes it possible to leverage *in silico* experiments to decipher sequence influences on intrinsic affinity and *in vivo* occupancy.

## Methods

### Yeast cell culture

The Myc-tagged Pho4 and Cbf1 *Saccharomyces cerevisiae* strains were obtained from Rick Young’s lab (113). For ChIP-nexus, cells were grown in 50 ml of yeast peptone dextrose (YPD) media at 25°C to OD_600_ ∼ 0.8. For Pho4 induction, cells were washed twice with and incubated at 25°C for 2h with 50 ml of phosphate-free media (YNB powdered media without phosphates from Fisher Scientific (MP114027812), supplemented with 2% glucose and 0.1g/L of sodium chloride).

### Yeast chromatin preparation

The yeast chromatin preparation was performed as previously described with minor modifications (52). For each experiment, 50 ml of cells were cross-linked at a final concentration of 1% formaldehyde (Fisher Scientific) for 15 min at 25°C. The reaction was quenched with 125 mM glycine for 5 min. Cells were washed three times with cold wash buffer (10 mM Tris-HCl, pH 7.5, 100 mM NaCl) at 4°C. After the last wash, the pellet was snap frozen in liquid nitrogen. The pellet was then resuspended in 700 µl of FA lysis buffer (50 mM HEPES-KOH pH 7.5, 150 mM NaCl, 1 mM EDTA, 1% triton-X-100, 0.1% Sodium deoxycholate, supplemented with protease inhibitor) and transferred to a 1.5 ml Sarstedt micro tube. 1 ml of 0.5 mm zirconia/silica beads was added to the tube and cells were lysed in a Mini-BeadBeater 24 (BioSpec products) for three cycles (3 min ON/5 min OFF). During the OFF cycle, samples were kept on ice. Cell lysates were separated from silica beads and collected in a new 1.5 ml tube using a 23G 1” needle (BD Microlance) and centrifuged at maximum speed for 4 min at 4°C to pellet the chromatin. The supernatant was discarded and the pellet was resuspended in 750 µl of FA lysis buffer supplemented with 0.1% SDS. Samples were sonicated with a Bioruptor Pico (Diagenode) for 18 cycles of 30 sec ON and 30 sec OFF intervals. Chromatin extracts were centrifuged at 16,000 g for 10 min at 4°C and the supernatant was used for the ChIP reaction.

### Antibodies

For each ChIP, 50 µl of protein G Dynabeads (Invitrogen) was washed three times with 1 ml of FA lysis buffer and after the last wash, resuspended in 500 µl of FA lysis buffer. 10 µg of anti-myc tag antibody (abcam 9E10) was added to the beads and incubated for 2 hours at 4°C with rotation. Following the incubation, antibody-bound beads were washed three times with 1 ml of FA lysis buffer.

### ChIP-nexus

Chromatin extract was added to the antibody-bound beads and incubated overnight at 4°C with rotation. The ChIP-nexus process was performed as previously described (37), with some modifications. Briefly, the chromatin was washed with Nexus wash buffers A to D (wash buffer A: 10 mM Tris-EDTA, 0.1% Triton X-100; wash buffer B: 150 mM NaCl, 20 mM Tris-HCl, pH 8.0, 5 mM EDTA, 5.2% sucrose, 1.0% Triton X-100, 0.2% SDS; wash buffer C: 250 mM NaCl, 5 mM Tris-HCl, pH 8.0, 25 mM HEPES, 0.5% Triton X-100, 0.05% sodium deoxycholate, 0.5 mM EDTA; wash buffer D: 250 mM LiCl, 0.5% IGEPAL CA-630, 10 mM Tris-HCl, pH 8.0, 0.5% sodium deoxycholate, 10 mM EDTA) and 10 mM Tris, pH 7.5. End-repair and dA-tailing enzymatic reactions were performed using the NEBNext End Repair and NEBNext dA-tailing module. The four fixed barcodes (ACTG, CTGA, GACT and TGAC) within ChIP nexus adapter mix were ligated with quick T4 DNA ligase. Following barcode extension, samples were treated with lambda exonuclease to generate high-resolution transcription factor binding footprints. Following reverse crosslinking and ethanol precipitation of DNA, purified DNA fragments were circularized and directly subjected to PCR amplification. The ChIP-nexus libraries were then gel purified before sequencing. For each factor, at least two biological replicates were prepared side-by-side from independent colonies.

### Library sequencing and data processing

ChIP-nexus libraries were sequenced in single-end mode (1 X 75 cycles) on a NextSeq 500 instrument. The adapters were trimmed using cutadapt (v 1.8.1)(114) and reads were aligned to the Saccharomyces cerevisiae (sacCer3) genome using bowtie (v1.1.2)(115). Reads were filtered using SAMtools (v1.3.1)(116) to remove unmapped reads and PCR duplicates. Reads aligned to the same position with the same barcode, CIGAR string and the SAM flag were deduplicated using nimnexus dedup (v.0.1.1)(46). The final filtered BAM file was converted to tagAlign format (BED 3 + 3) using bedtools ‘bamtobed‘ (v.2.26)(117). Cross-correlation scores were obtained for each file using phantompeakqualtools (v.1.2)(118). BigWig tracks containing the strand-specific number of aligned 5′ read ends (pooled across all replicates) were generated using bedtools genomecov –5 –bg –strand <+/−>, followed by bedGraph to BigWig conversion using UCSC bedGraphToBigWig v.4. Peaks were called using MACS2 (v.2.1.1)(119) by extending the 5’ ends of reads on each strand using a 150-bp window and then computing coverage of extended reads across both strands (shift = –75, extsize = 150). For each TF, peak calling was performed on filtered, aligned reads from each replicate using a relaxed threshold P=0.1. The ChIP–nexus pipeline performing the described steps (for example, turning raw reads in FASTQ format to BigWig coverage tracks and called peaks) is available at https://github.com/mlweilert/chipnexus-processing-scripts and https://github.com/kundajelab/chip-nexus-pipeline.

### PB-exo

All the sequencing files and peak files for Pho4 and Cbf1 as well as the control (input) replicates were downloaded from the NCBI Gene Expression Omnibus (GEO; https://www.ncbi.nlm.nih.gov/geo/) under accession number GSE93662 (52). The datasets were processed using the ENCODE pipeline: https://github.com/ENCODE-DCC/chip-seq-pipeline2 (version: v1.3.6) with fraglen set to 150 and shift set to –75.

### ChIP-seq

All the sequencing files for human TFs were downloaded from the ENCODE portal (41) and processed using the ENCODE pipeline: https://github.com/ENCODE-DCC/chip-seq-pipeline2 (version: v1.3.6). For the Glucocorticoid receptor data, we used treatment time series ENCSR210PYP at the two hours time point ENCSR720DXT. For Myc and MAX in HELA-S3, we used the control run ENCSR000EZM and the experiments ENCSR000EZD and ENCSR000EZF, respectively. For Myc in K562, we used the experiment ENCSR000EGS and the control run ENCSR000FBB. For GABPA in Hep-G2, we used the experiment ENCSR000BJK and the control run ENCSR000BLH. For GABPA in liver tissue, we used the experiment ENCSR038GMB and the control run ENCSR019XRC. For E2F1 in MCF7, we used the experiment ENCSR000EWX and the control run ENCSR000EWW. For MAX in NB4, we used the experiment ENCSR000EHS and the control run ENCSR000FAM.

### Neural network training and evaluation

#### Model architecture

We used modified versions of the BPNet architecture. BPNet is a sequence-to-profile convolutional neural network that uses one-hot-encoded DNA sequence as input to predict base-resolution read count profiles as output. The output of the final convolutional layer (or the bottleneck activation map) serves as input for two output heads: (1) a deconvolutional layer for the profile prediction; and (2) a global average pooling layer followed by the fully connected layer for total read count prediction. When available, control tracks are provided to the model. The full details of BPNet as well as its implementation are publicly available (46).

For the PB-exo models, the input sequences were 546 bp. The first convolutional layer used 64 filters of width 21 bp, followed by 6 dilated convolutional layers (each with 64 filters of width 3). The deconvolutional layer had a filter width of 75 bp and the output profile length was 200 bp.

The ChIP-nexus models had the same number of layers, but with 25 filters in each layer instead of 64. The deconvolutional layer had a filter width of 25 bp, resulting in output profile length of 250 bp.

All the mammalian ChIP-seq models had the same architecture. The input sequences were 1346 bp. The first convolutional layer used 64 filters of width 21 bp, followed by 6 dilated convolutional layers (each with 64 filters of width 3). The deconvolutional layer had a filter width of 75 bp and the output profile length was 1000 bp.

For all models, we set the count prediction head relative weight compared to the profile prediction head to 100. All the convolutional and deconvolutional layers used in our models were reverse-complement layers (120).

#### Model training

For the PBexo models, the reported peaks (52) were divided for training, validation, and testing. For Cbf1, regions from chromosomes III and XV (106 regions, 10%) were used as the tuning set for hyperparameter tuning. Regions from chromosomes V and VI (105 regions, 10%) were used as the test set for performance evaluation. The remaining regions (846 regions, 80%) were used for model training. For Pho4, regions from chromosomes V and IX (576 regions, 10%) were used as the validation set. Regions from chromosomes I and XII (552 regions, 10%) were used as the test set. The remaining regions (4558 regions, 80%) were used for model training.

For the ChIP-nexus models, overlap peaks were divided for training, validation, and testing. For Cbf1, regions from chromosome VII (118 regions, 10%) were used as the tuning set for hyperparameter tuning. Regions from chromosomes IX and XI (117 regions, 10%) were used as the test set for performance evaluation. The remaining regions (939 regions, 80%) were used for model training. For Pho4, regions from chromosome XV (162 regions, 10%) were used as the validation set. Regions from chromosomes VIII and X (162 regions, 10%) were used as the test set. The remaining regions (1297 regions, 80%) were used for model training.

For human ChIP-seq models, IDR peaks were divided for training, validation, and testing. Regions from chromosomes 8 and 9 (10%) were used as the test set for performance evaluation. Regions from chromosomes 16, 17, and 18 (10%) were used as the tuning set for hyperparameter tuning. The remaining regions were used for model training.

Outside of these particular splits, cross validation was performed in five folds, achieving similar performance across folds (See Supplement). Hyperparameters were adjusted to yield best performance on the tuning set. During training, we deployed a uniform jitter with a maximum shift of 200 bp of the regions in each batch. All neural network models were implemented and trained in Keras (v.2.2.4) (TensorFlow backend v.1.14)(121,122) using the Adam optimizer (123) with learning rate 0.001 and early stopping with patience of ten epochs.

#### Model cross-validation

In addition to the aforementioned training, all models were evaluated in five fold cross-validation (See supplemental materials). Each fold divided the genome into contiguous non-overlapping regions with at least a 2 Kbp buffer between the end of one set and the beginning of another. The folds always assigned 10% of peaks to the testing set and 10% of peaks to the validation set. The remaining peaks were assigned to the training set.

#### DeepSHAP contribution scores

We used the deep explainer implementation of SHAP (124), which is an updated version of the DeepLIFT algorithm (125), to interpret all models. We used a shuffled reference with 20 random shuffles. At each position in the input sequence, we iterated over the one-hot encoding possibilities and computed the hypothetical difference-from-reference in each case. We then multiplied the hypothetical differences-from-reference with the multipliers to get the hypothetical contributions.

For each of the one-hot encoding possibilities, the hypothetical contributions were then summed across the ACGT axis to estimate the total hypothetical contribution of each position. This per-position hypothetical contribution was then projected onto whichever base was present in the hypothetical sequence. The reason this is a fast estimate is that the multipliers are computed once using the original sequence, and are not computed again for each hypothetical sequence.

#### GC-matched augmentation

For the GC-augmented models, the GC content of each peak region was computed. Then the negative set, which is at least 1Kbps from any peak, was searched for a window that most closely matches the GC content of each peak region. Jittering the positive regions ensured a positive:negative ratio of 10:1. Finally, the augmented dataset was divided by chromosome in a way that ensured no overlap between training, validation, and testing.

### Affinity Distillation

#### In-silico marginalization

In silico marginalization relies on the counts head of BPNet. To obtain a marginalization score for a sequence: (1) Background sequences are generated by dinucleotide shuffling DNA sequences from held-out genomic peaks, (2) the sequence of interest is inserted at the center of the background sequences, (3) the model predictions from the count head are stored for both the background sequences and the sequences with the insert, (4) the mean of the differences between the two sets of predictions (mean predicted log count ratio) across the different backgrounds (Δ log(counts)) is the marginalization score for the sequence of interest.

#### Calibration

Calibration performs inference of the binding free energies using a regression model. Using a sample of *in vitro* measurements, we fit a linear regression model with the generated marginalization scores as the input. The resulting function recovers the inference we would have obtained if the neural network model was predicting in the relative free energy space. We can generalize to other sequences, without access to measurements, by deploying a correction for inference using the fitted regression model.

### Motif discovery methods

*STREME*. STREME from MEME suite (v.5.4.1)(126) was run on the central 100bp around summits of peaks. Prior reports confirm that using the central 100 bp works extremely well with ChIP-seq for motif discovery (127). Shuffled set was used as the control.

*Weeder2.* Weeder 2.0 (128) was run on the central 100bp around summits of peaks. For yeast datasets, all peak regions were used. For human TFs, Weeder2 was run on a sample of 1000 regions (flag –top 1000). All runs were performed using ChIPseq mode (flag –chipseq).

*TF–MoDISco.* TF–MoDISco–Lite (v2.0.7)(63) was run on counts contribution scores for each TF (using all peak regions bound by the TF on autosomes) with sliding windows of size 20 bp, flanks of size 5bp, the false-discovery rate threshold of 0.05 (target_seqlet_fdr), and a maximum of 20,000 seqlets per metacluster. All CWMs were trimmed down to their final lengths by removing flanking positions using the default trimming function and threshold. For all discovered motifs, CWMs were computed from the aligned seqlets by averaging the base frequencies and contribution scores, respectively. All the results presented in this study also held when using TF–MoDISco (v.0.5.5.5), which runs a different sorting algorithm.

### Methods Comparison

To quantitatively assess the performance of different methods as well as Affinity Distillation, we used each method to compute the affinities across all sequences in the given libraries. For each motif discovery method, we used all the returned motifs, up to the top ten motifs. For motif scoring, both the sequence and its reverse complement were scored and the higher score was chosen (Supplement **Section 7**). The scores for each method, including Affinity Distillation, were calibrated using the same calibration set. Finally, we calculated and reported the post calibration RMSE for each method.

### gcPBM

For Pho4 and Cbf1 gcPBM (29), the fluorescent intensities (Alexa 488 Adjusted) were downloaded from GEO, accession GSE163512. We took the natural log of the intensities. For multiple measurements of the same sequence, we computed the mean. For Pho4, the 400nM set of measurements was used, and the 1µM set was used for Cbf1.

For human TFs, the fluorescent intensities were downloaded as log normalized gcPBM signals (30) from GEO, accession number GSE97794. The 36-mer gcPBM probes (both putative binding sites and negative controls), their genomic coordinates, and the associated measurements were downloaded from the same study. For GABPA, the 100nM set of measurements was used, and the 250nM set was used for E2F1.

### Differential Specificity Methods

We compared gene expression data from (68) downloaded from GEO, accession number GSE23580. Comparing wild type no Pi versus high Pi conditions identified Pho4 genes under physiological conditions (first set of genes). Comparing ΔPho80ΔCbf1 versus ΔPho80ΔPho4ΔCbf1 in high Pi conditions identified the influence of Cbf1 on the gene activation role of Pho4 (second set of genes). Then, we downloaded the gcPBM signals from (29) for the Pho4 binding sites potentially responsible for the regulation of each target gene using the reported criteria: 1) the site was located within 1000bp upstream of the gene TSS, and 2) the in vitro Pho4-DNA binding level at the site was higher than the binding level at any of the negative control probes in the PBM experiment. We used the PBM intensities to compute the observed differential signal for each of the two sets of genes.

### GR MITOMI Methods

#### Fabrication of microfluidic molds and devices

Flow and control molds were fabricated as described previously (86)and all design files are available on the Fordyce Lab website (http://www.fordycelab.com/microfluidic-design-files). We fabricated two-layer MITOMI devices from these molds using polydimethylsiloxane (PDMS) polymer (RS Hughes, RTV615). The control layer was fabricated with a 1:5 ratio of cross-linker to base and the flow layer was fabricated with a 1:20 ratio of cross-linker to base. Layers were aligned manually and holes were punched with a mounted catheter hole punch.

#### Preparation of MITOMI oligonucleotide arrays

**Pseudorandom 8mer libraries were designed using the ShortCAKE algorithm**(85). Single-stranded DNA (ssDNA) oligonucleotides were synthesized by Integrated DNA Technologies (IDT). Oligos were fluorescently labeled and converted to double-stranded DNA (dsDNA) in a 15 μL Klenow extension reaction:

- dNTPs (final conc: 100 μM each)
- Cy5-labeled primer (final conc: 10 μM)
- oligonucleotide (final conc: 10 μM)
- 2 units, Klenow fragment (exo-) (NEB)
- NEBuffer 2 (final conc: 1x)

All reagents were mixed together except for the Klenow enzyme. Samples were denatured at 94°C for 3 min and cooled to 37°C over 45 minutes. Upon reaching 37°C, Klenow enzyme in 1x NEBuffer 2 was added to the sample. Oligonucleotides were then allowed to extend for 1 hour at 37°C and the reaction was heat-inactivated at 72°C for 20 min. Primer extension was confirmed by denaturing polyacrylamide gel electrophoresis.

DNA reactions were mixed with PEG (MW=6,000, Fluka) and trehalose (Fluka) to final concentrations of 0.125% PEG and 12.5 mg/mL trehalose in 3x SSC to promote solubilization of printed DNA. For the pseudorandom 8mer library, DNA was printed at a single concentration; for subsequent libraries, reactions were serially diluted to produce 8 different concentrations. This library of serial dilutions was then spotted onto 2”x3” epoxysilane-coated glass slides (Thermo Scientific, UCSF2X3-C50-20) with a UCSF/Stanford-style custom microarrayer with 75-μm silicon tips (Parallel Synthesis). After printing, we manually aligned MITOMI devices to the DNA array such that each chamber contained one oligonucleotide spot. Devices were then bonded to the glass slide by baking on a hot plate at 80°C for 4 hours to overnight.

#### Transcription factor expression for MITOMI

Linear templates for in vitro transcription and translation of full-length GR isoform alpha (NCBI Reference Sequence NP_000167.1) were generated by a series of 2 PCR reactions as described previously (86). The first reaction adds a Kozak sequence, C-terminal 6x His tag and a stop codon to the open reading frame of the glucocorticoid receptor gene. The second reaction amplifies this product and adds a T7 promoter, a –globin spacer to improve expression, a poly-A tail, and a T7 terminator. PCR reactions were cleaned with a generic column cleanup protocol and then TOPO cloned (Invitrogen) into the pCR-2.1-TOPO vector prior to IVTT.

Proteins were expressed in rabbit reticulocyte lysate (TNT T7 Quick Coupled Transcription/-Translation System, Promega) according to the manufacturer’s instructions. 75 μL reactions were prepared for each device with 1 μg of PCR product. Fluorotect Green BODIPY-FL-labeled charged lysine tRNA (Promega) was added to allow protein visualization. Reactions were incubated at 30◦C for 90-120 minutes while being shaken at 600 rpm.

#### MITOMI device operation and experimental pipeline

Devices were run as described previously (86), with flow and control pressures at 5 and 12-20 psi, respectively. The slide surface was functionalized with biotinylated BSA (bBSA; ThermoFisher, 2 mg/mL) for 20-30 minutes and then washed for 5-10 minutes with 50 mM HEPES (pH 8.0). NeutrAvidin (ThermoFisher, 1 mg/mL) was then flowed over the device for 20-30 minutes and the device was washed again with HEPES. Button valves were then pressurized and the remaining surface was passivated with bBSA, flowed for 30 minutes followed by a 10-minute HEPES wash. Button valves were opened and biotinylated anti-pentaHis antibody (Qiagen, 1:4 dilution in HEPES) was flowed for 30 minutes, followed by a 10 minute HEPES wash. Finally, IVTT-expressed His-tagged protein was flowed for 40 minutes.

The neck valves were then opened to allow for solubilization of printed DNA oligonucleotides. Simultaneously, sandwich valves were closed to spatially separate adjacent chambers and prevent mixing of DNA sequences. The TF-DNA binding reaction was allowed to proceed for 60 minutes before the button valves were closed. Sandwich valves were then opened and the device was washed with HEPES to flush unbound DNA. Slides were then scanned with an ArrayWorx scanner for 1 s at 488 nm to measure BODIPY-FL-labeled protein abundance and 5 s at 633 nm to measure Cy5-labeled DNA abundance.

#### MITOMI image processing

Array scans were analyzed with GenePix v6.0 software to quantify Cy5 and BODIPY-FL intensities under the button valve and Cy5 in the DNA chambers. Local background signal was subtracted to account for differences in local illumination, etc. All further analyses were performed with custom software written in Python. Chambers with unusually high or low BODIPY-FL signal and chambers with obvious debris or protein aggregates were omitted from further analyses.

#### Identification of preferred half-sites

Pseudorandom 8mer data were analyzed by computing normalized intensity ratios, running fREDUCE to identify globally preferred sequences, and then running MatrixREDUCE to refine motifs, as described previously (86).

#### Determination of equilibrium binding energies

For each chamber, we quantified the ratio (R) of button DNA fluorescence (Cy5) to button protein fluorescence (BODIPY-FL). We also quantified chamber DNA fluorescence, which is proportional to DNA concentration. We then collated concentration-dependent binding data by oligonucleotide sequence within each experiment and fit to a Langmuir isotherm:

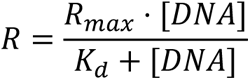

where *R_max_*, the saturation value, is globally fit for all data on a device (as in (22,86)) and where Kd is fit for each oligo sequence. Due to a lack of calibration data relating chamber fluorescence to [DNA], we report ΔΔG values, as they are relative and eliminate the need for calibration.

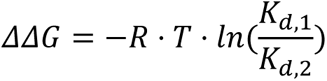

To account for fitting errors associated with a globally fit *R_max_* value, we iteratively selected half of the available chambers on a device (e.g. if 1,200 chambers were left after data QC, we selected 600 chambers without replacement) and performed a fit on this subset. We repeated this process 1,000 times to build a distribution of estimated *K*_*d*_ values per oligo per experiment. Reported Kd values and errors are the mean and standard deviations of the *K*_*d*_ distributions per oligo per experiment.

### Code and Data Availability

All the ChIP-nexus data generated and used for this study are available in GEO:GSE207001. All the MITOMI measurements of glucocorticoid receptor binding are available at Zenodo: https://zenodo.org/record/6762262. The code to reproduce the results of this manuscript is available at: https://github.com/kundajelab/affinity_distillation.

## Supporting information

Model performance supplementary table

## Acknowledgments

We thank Jin-Wook Lee for his assistance with processing GR ChIP-seq data. We thank Arjun Aditham for his feedback on this manuscript. A.M.A. was supported by the Stanford Bio-X Fellowship. All the figures were created with Biorender. This research was supported by an NSF CAREER award to P.M.F. and P.M.F. is a Chan Zuckerberg Biohub Investigator. This research was supported by National Institutes of Health (NIH) grant numbers U01HG009431, U01HG012069, U24HG012343 to A.K.

