## Supplementary Figures for "*De novo* distillation of thermodynamic affinity from deep learning regulatory sequence models of *in vivo* protein-DNA binding"

### Section 1: PSAM evaluations

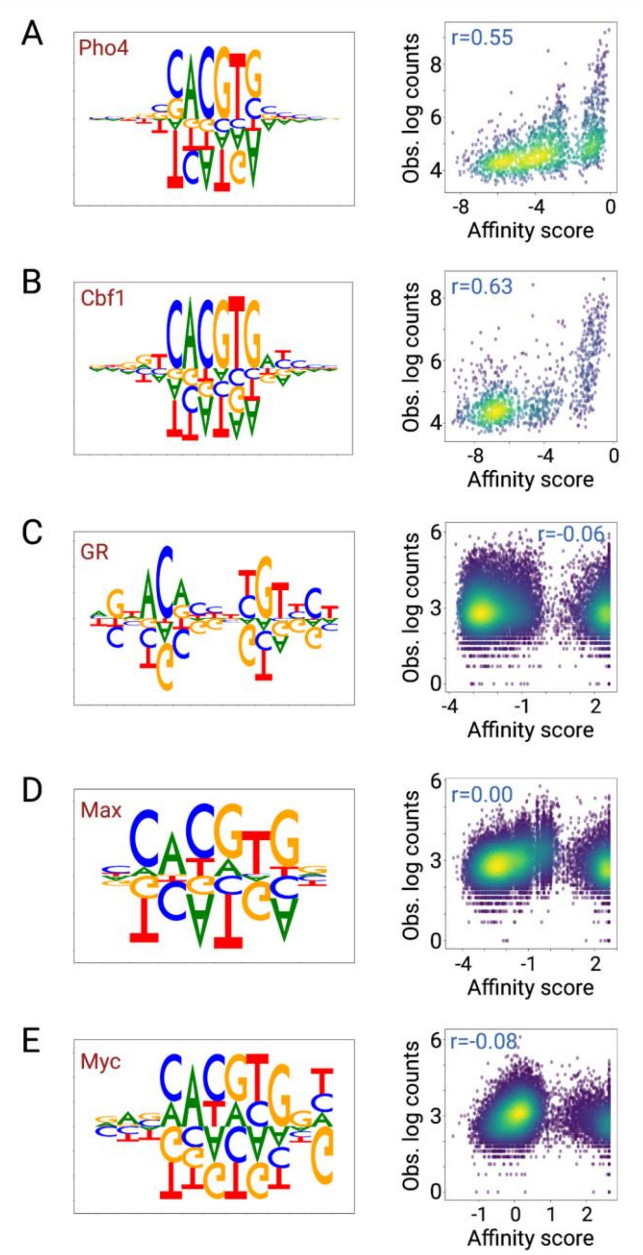


**Supplemental figure 1. Binding predictions based on *in vitro*-derived PSAMs. (A)** Pho4 PSAM derived from Maerkl & Quake data (1) and BET-seq experiments (2) for Pho4 in ΔΔG space (left) and comparison between PSAM-predicted affinities and observed log-transformed total counts in a 100bp window around genomic summits from ChIP-nexus data (right). **(B)** Same as A for Cbf1. **(C)** GR PSAM derived from MITOMI experiments in ΔΔG space (left) and comparison between PSAM-predicted affinities and observed log-transformed total counts in a 100bp window around genomic summits from ChIP-seq data (right). **(D)** Max normalized motif (3) (left) and comparison between PSAM-predicted affinities and observed log-transformed total counts in a 100bp window around genomic summits from ChIP-seq data from HeLa S3 cells (right). **(E)** Same as D for Myc.

**Section 2: MoDISco motifs**

One of the challenges associated with extracting affinities from neural networks is that the models learn many different motifs to achieve good accuracy. The strength of AffinityDistillation is that it side-steps the motif selection problem by directly scoring sequences of interest. In this section, we show the diversity of motifs learned by BPNet models. For each model, we show all the learned MoDISco motifs(4), generated by MoDISco–Lite. We report the task, pattern rank, and number of seqlets associated with each motif.

**Pho4 PBexo model**

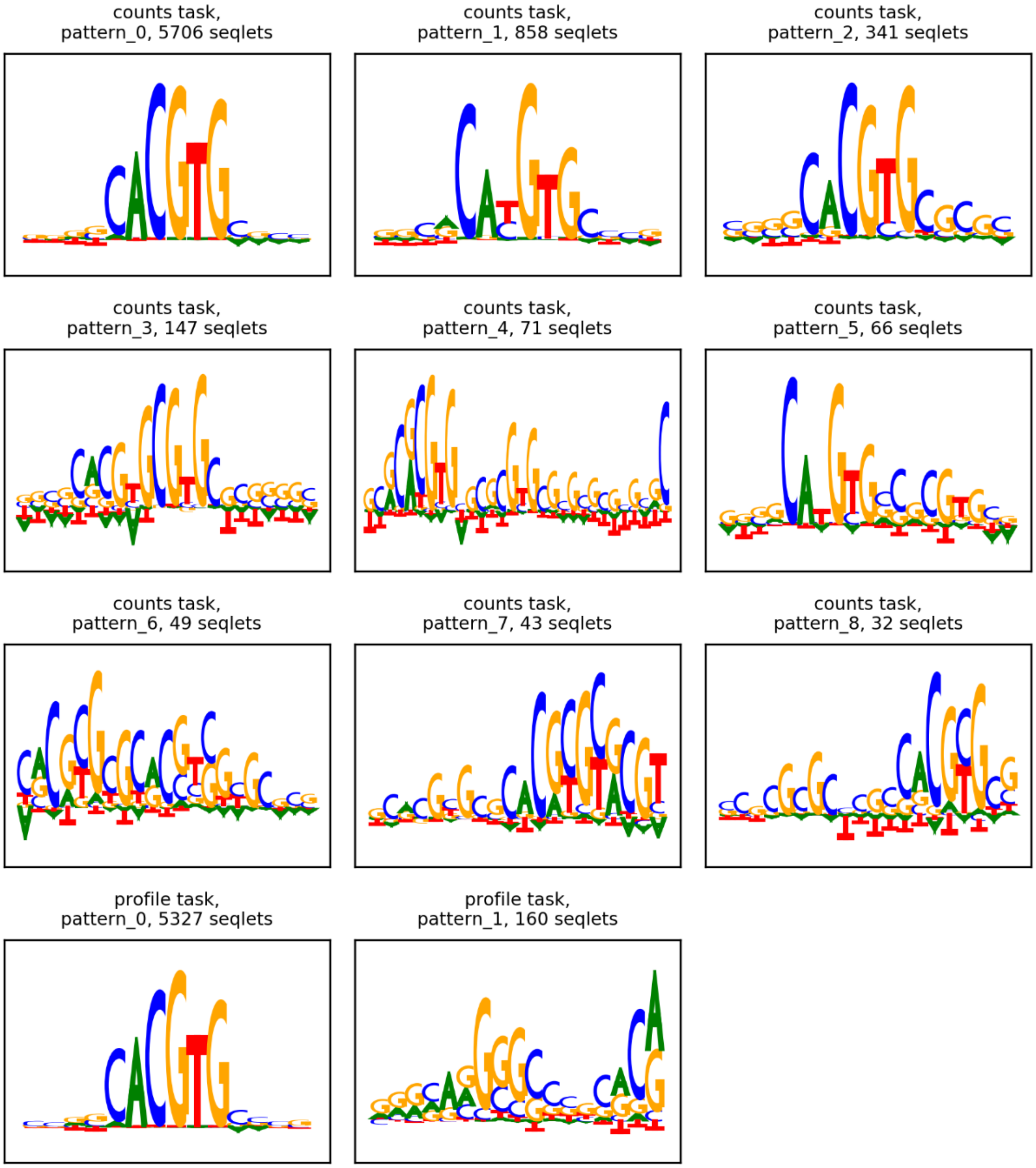

### Cbf1 PBexo model

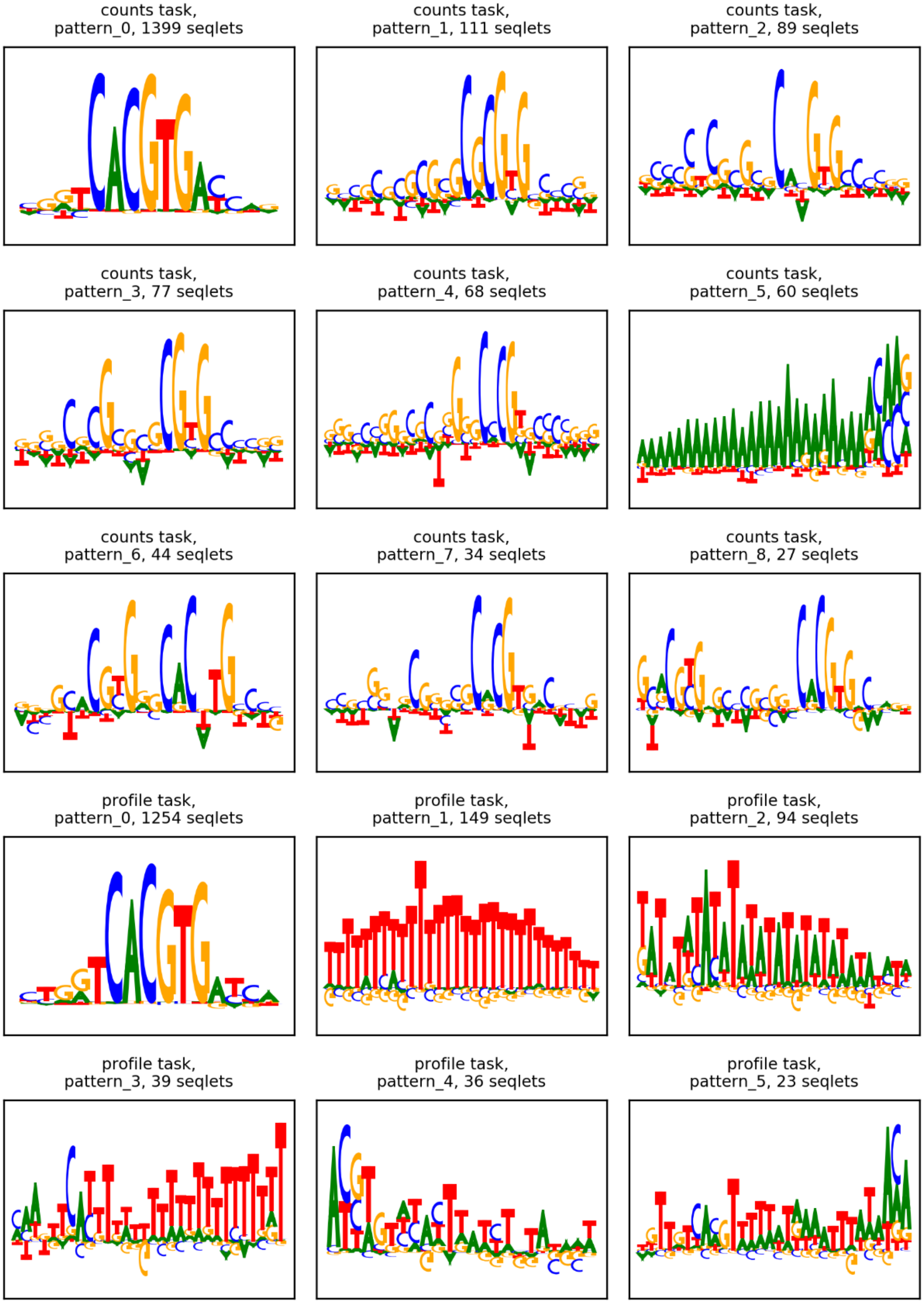

**Pho4 ChIP nexus model**

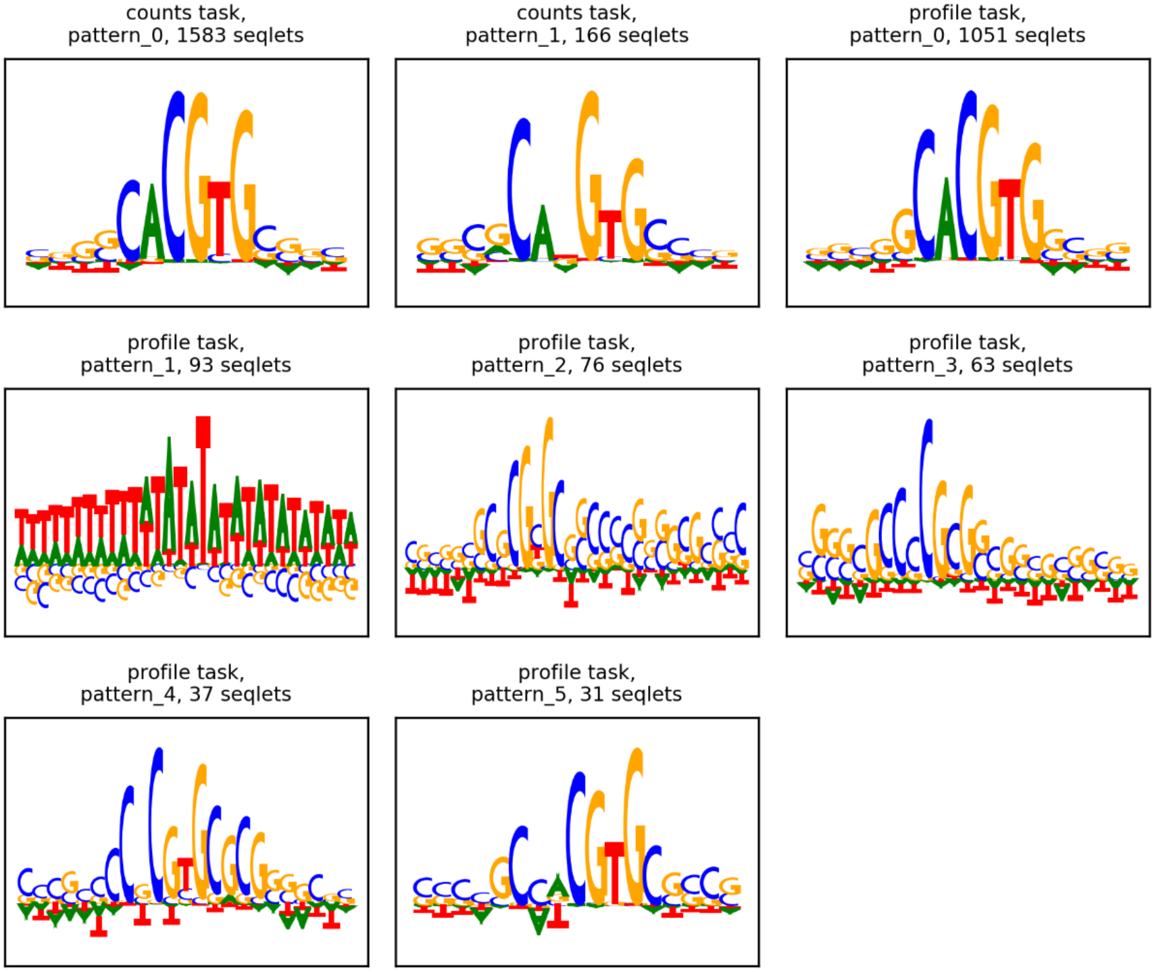

**Cbf1 ChIP nexus model**

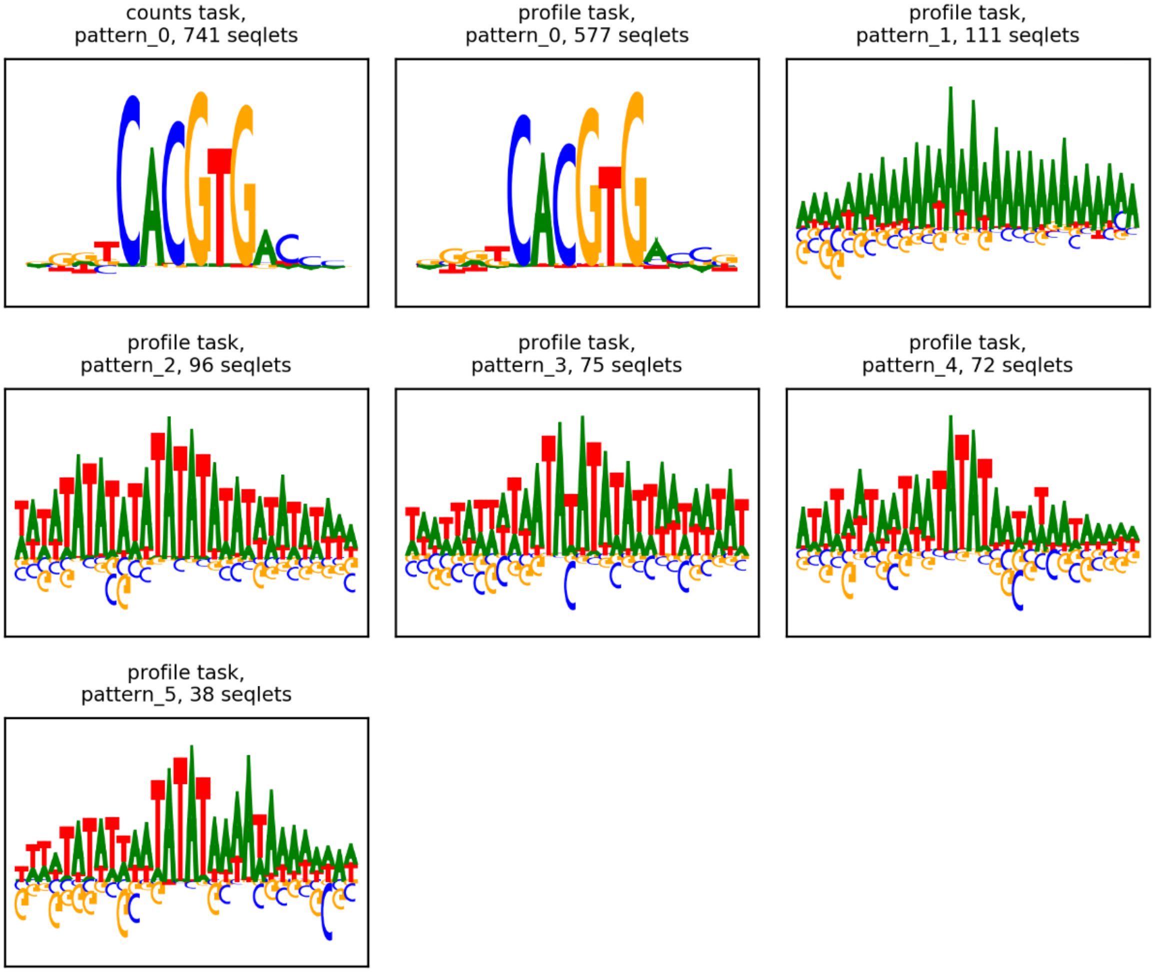


**MAX in HeLa-S3 ChIP seq model**

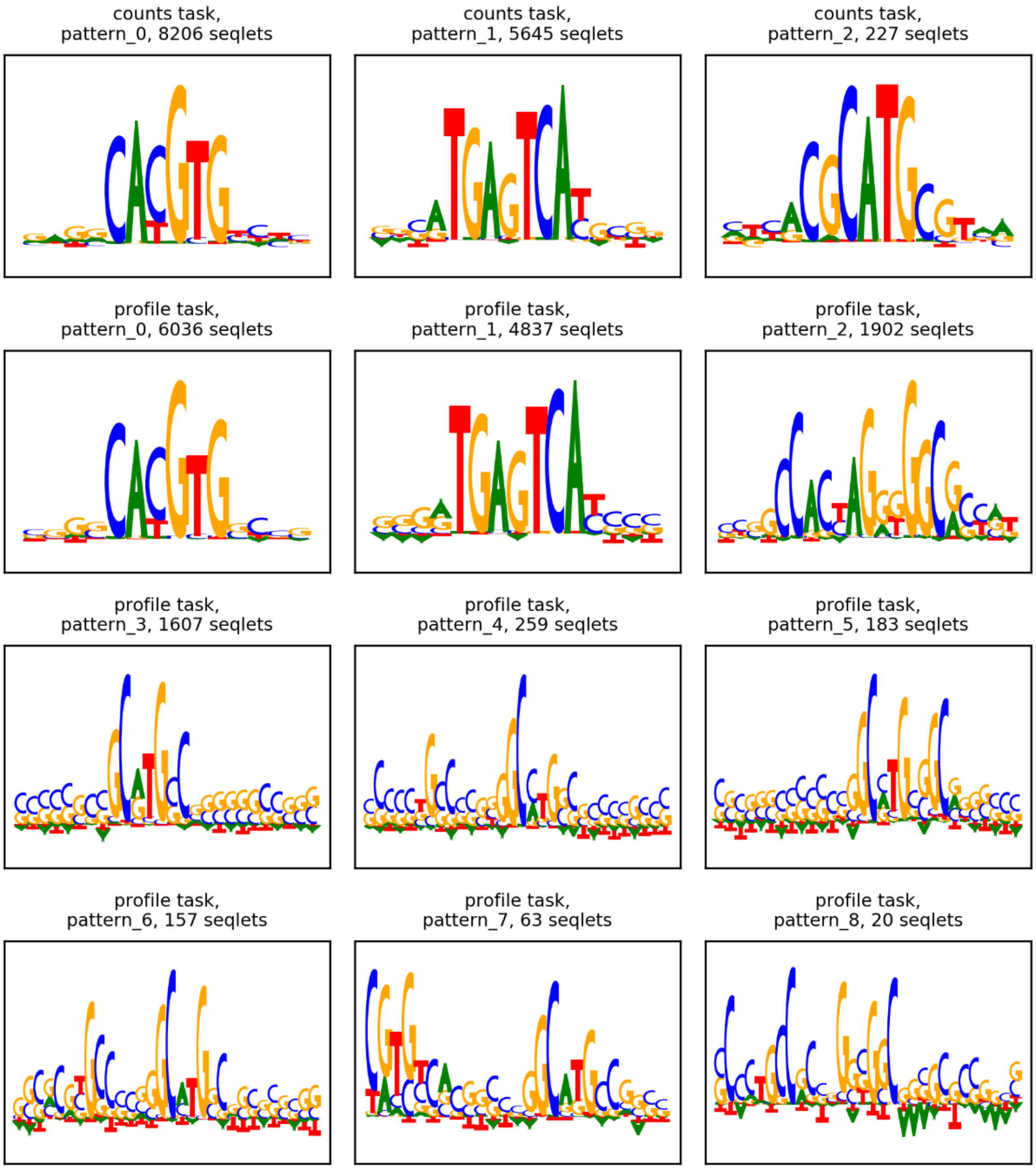


**Myc in HeLa-S3 ChIP seq model**

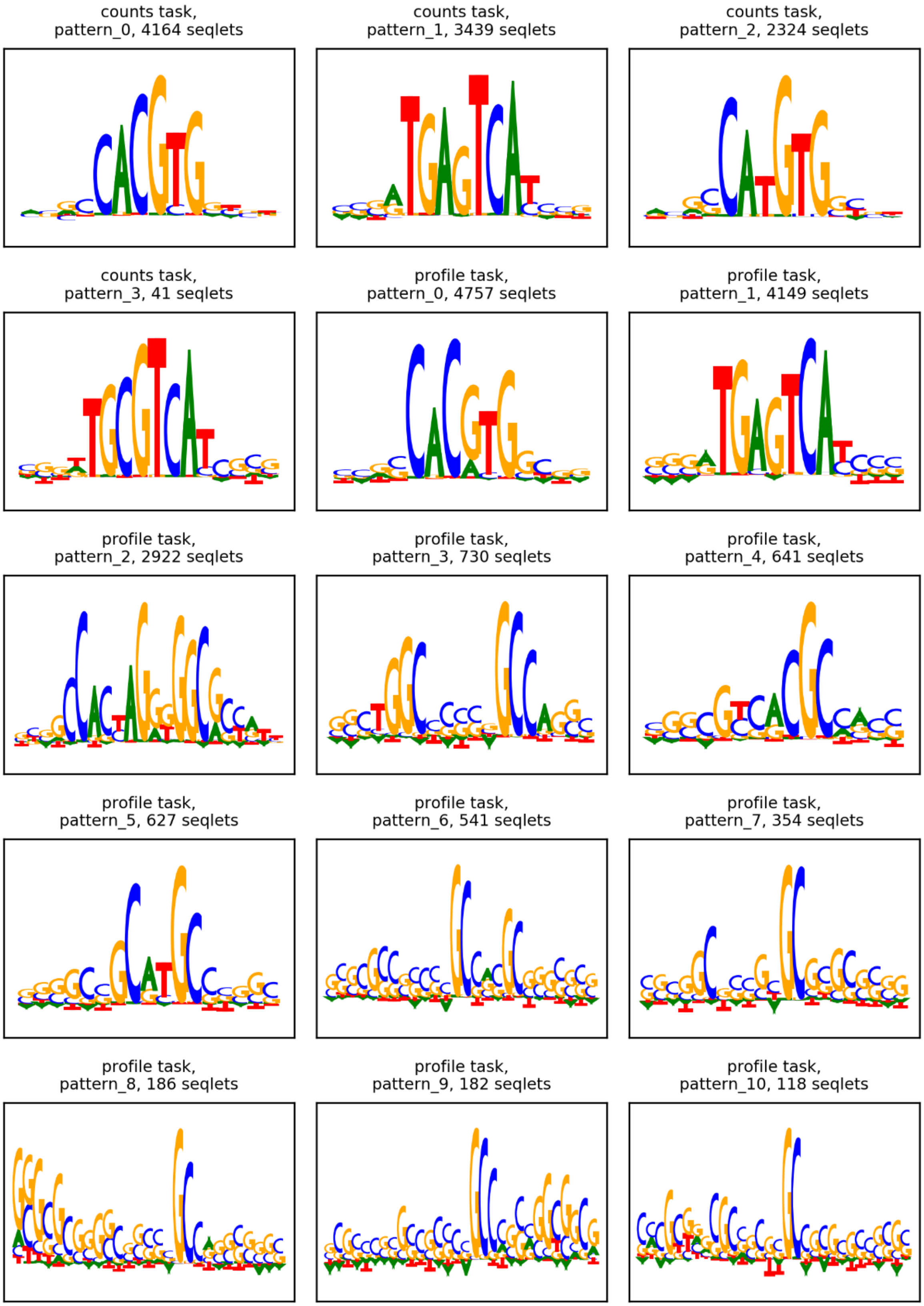


**Myc in K562 ChIP seq model**

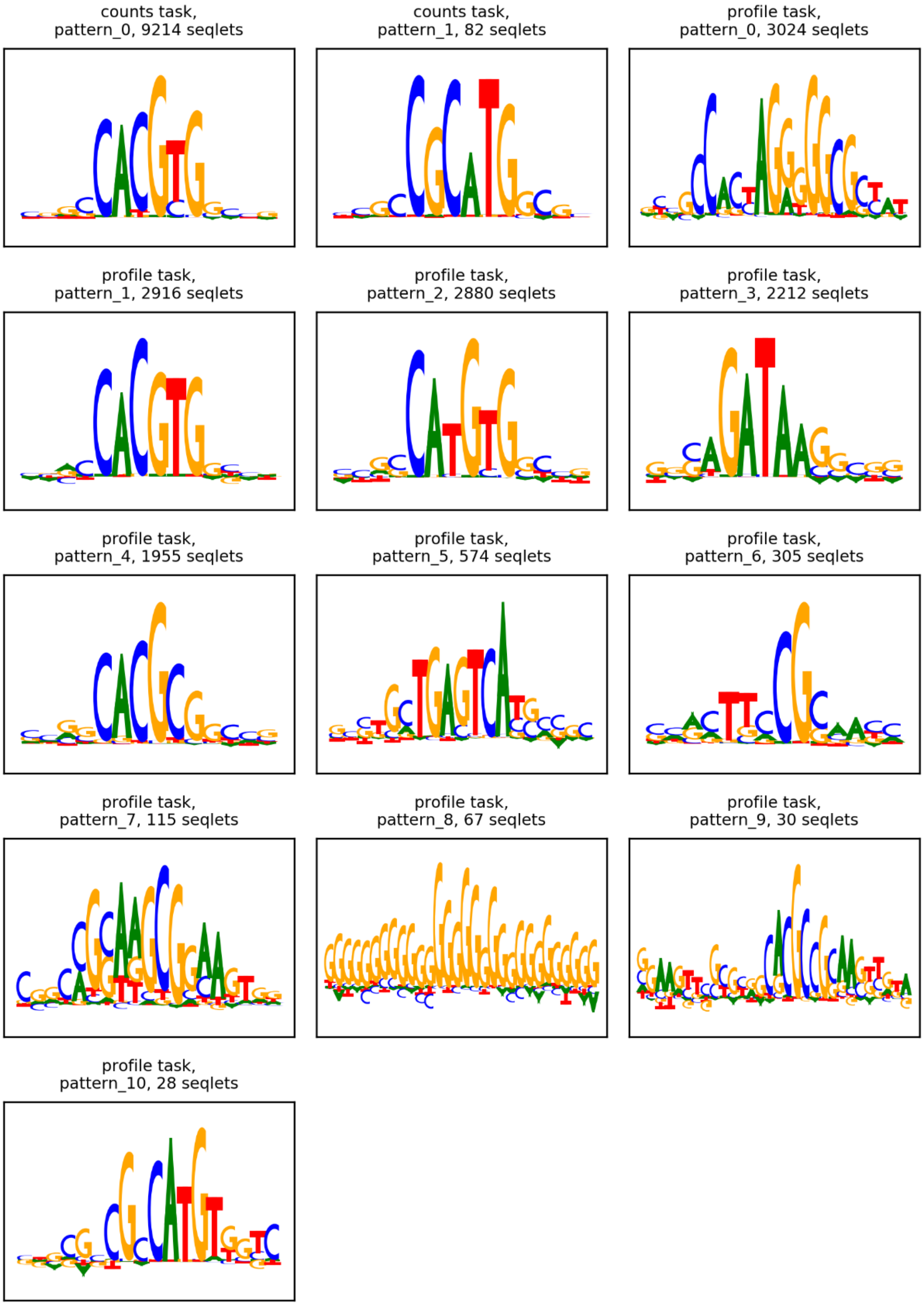


**GABPA in HepG2 ChIP seq model**

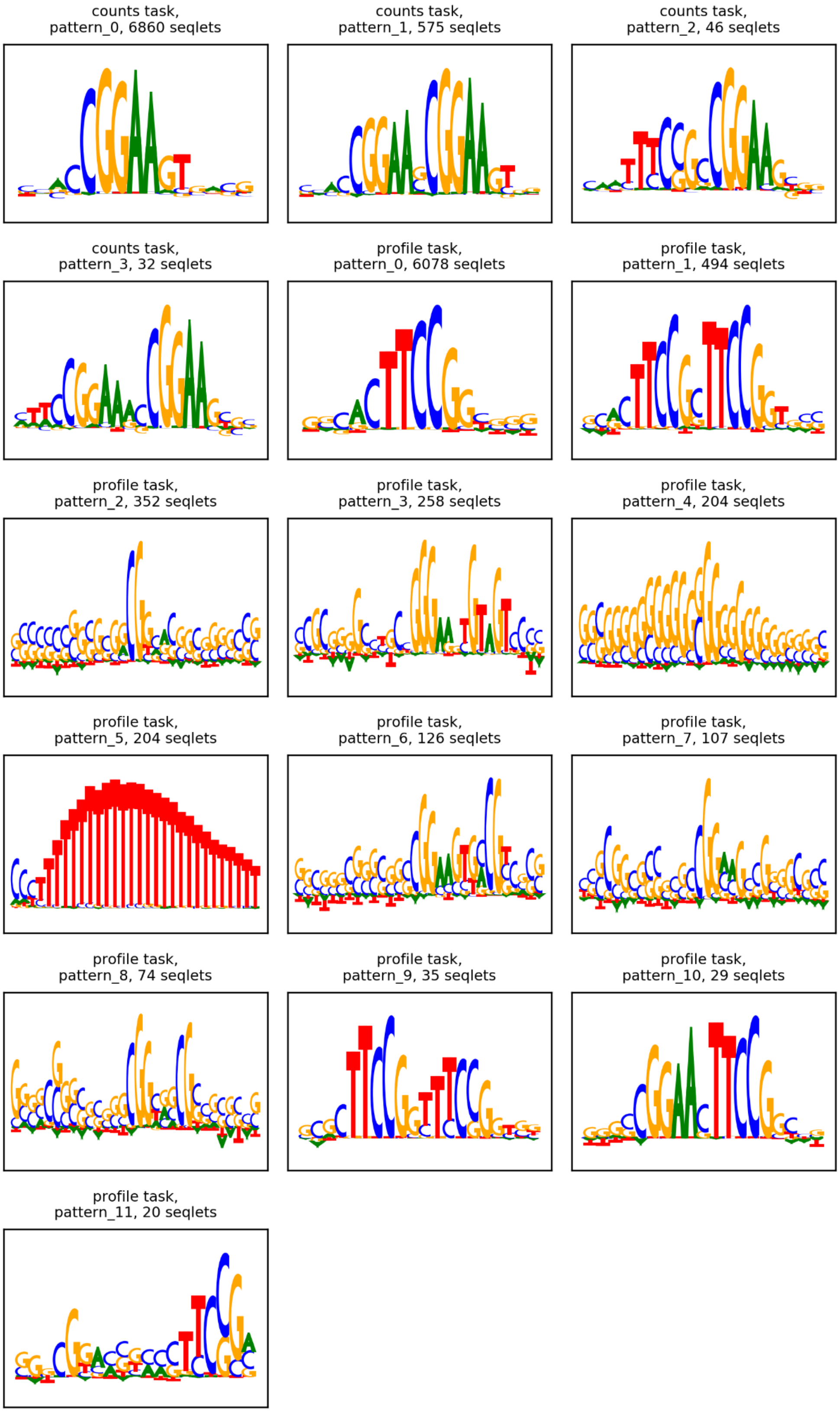


**GABPA in liver tissue ChIP seq model**

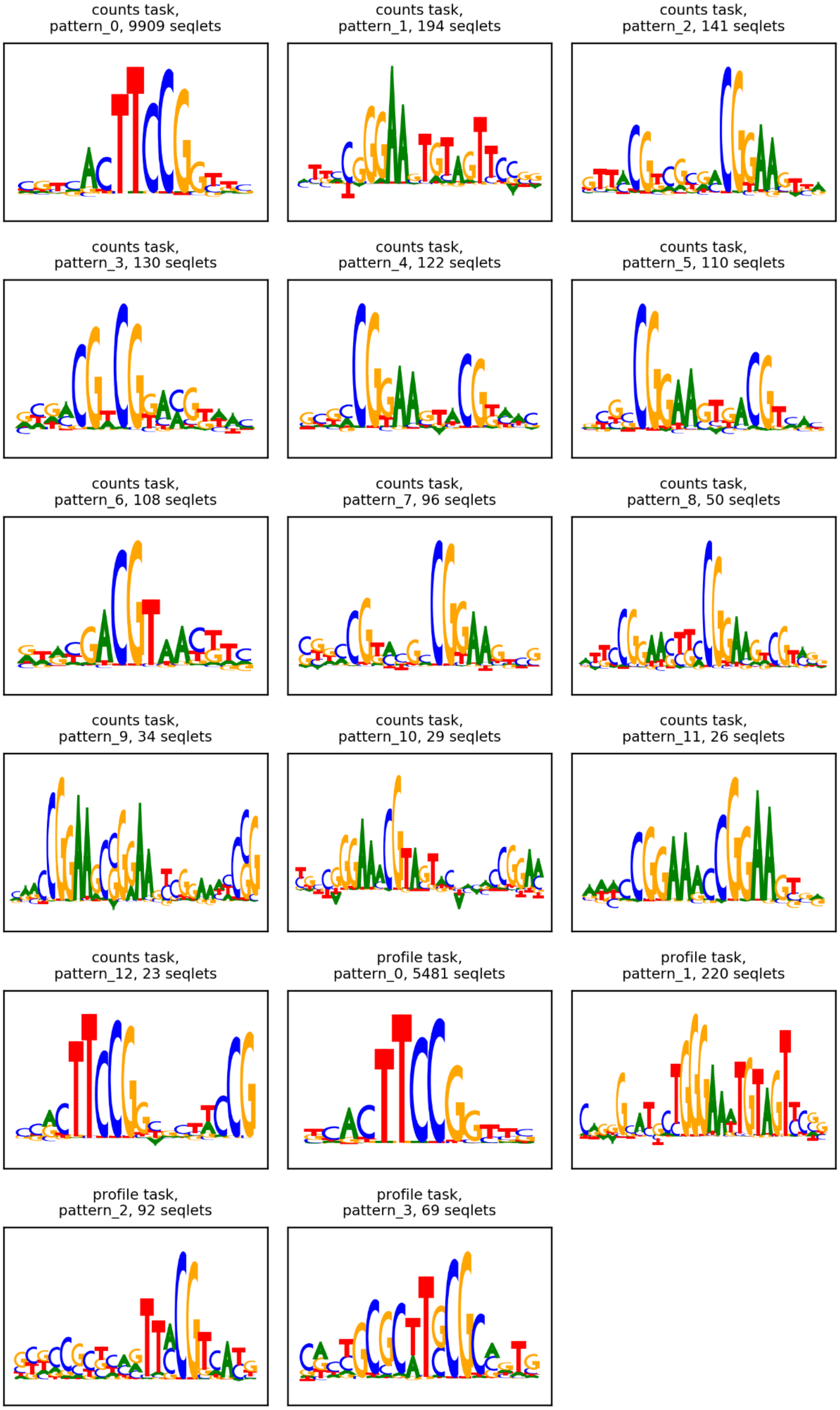


**E2F1 in MCF7 ChIP seq model**

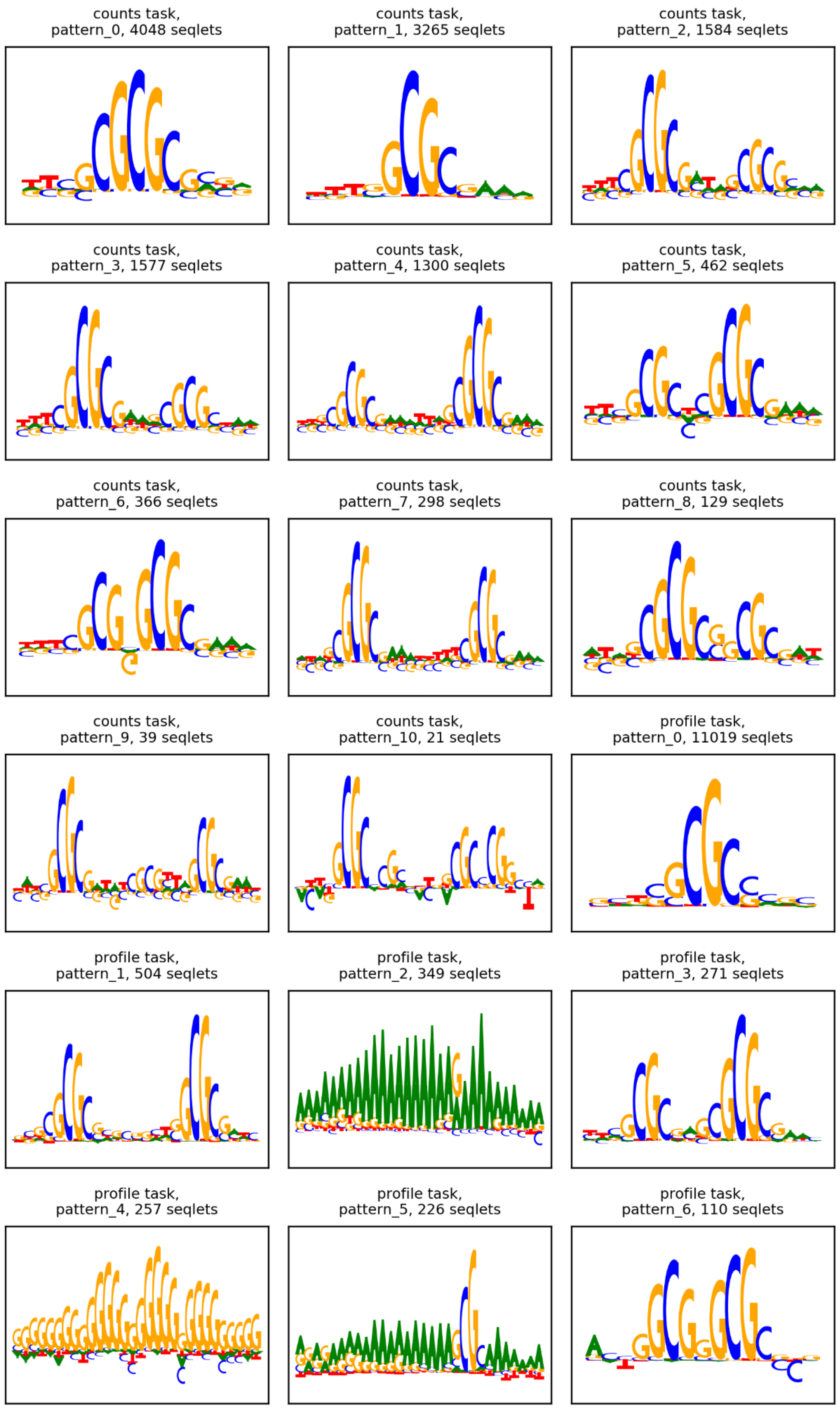
counts task,
pattern_0, 4048 seqlets
counts task,
pattern_1, 3265 seqlets
counts task,
pattern_2, 1584 seqlets
counts task,
pattern_3, 1577 seqlets
counts task,
pattern_4, 1300 seqlets
counts task,
pattern_5, 462 seqlets
counts task,
pattern_6, 366 seqlets
counts task,
pattern_7, 298 seqlets
counts task,
pattern_8, 129 seqlets
counts task,
pattern_9, 39 seqlets
counts task,
pattern_10, 21 seqlets
profile task,
pattern_0, 11019 seqlets
profile task,
pattern_1, 504 seqlets
profile task,
pattern_2, 349 seqlets
profile task,
pattern_3, 271 seqlets
profile task,
pattern_4, 257 seqlets
profile task,
pattern_5, 226 seqlets
profile task,
pattern_6, 110 seqlets

profile task,
pattern_7, 86 seqlets

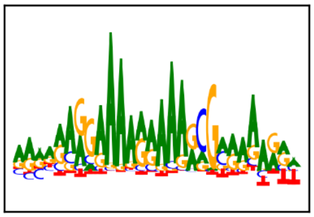

profile task,
pattern_8, 80 seqlets

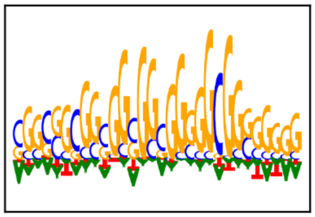

profile task,
pattern_9, 65 seqlets

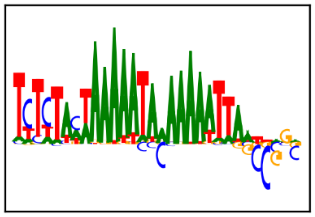

profile task,
pattern_10, 42 seqlets

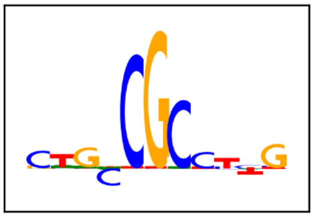

profile task,
pattern_11, 39 seqlets

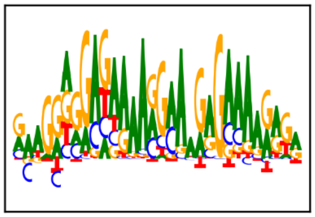

profile task,
pattern_12, 34 seqlets

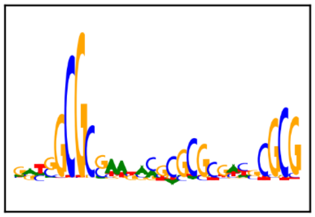

profile task,
pattern_13, 33 seqlets

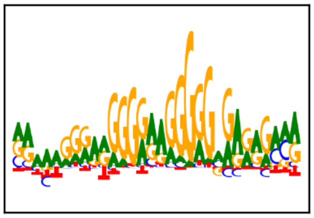

profile task,
pattern_14, 26 seqlets

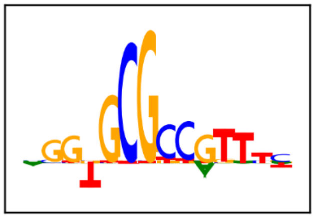

profile task,
pattern_15, 23 seqlets

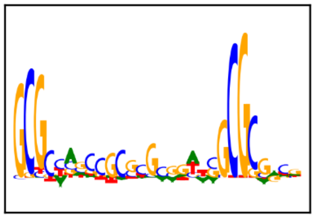

profile task,
pattern_16, 22 seqlets

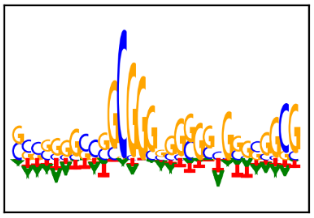

profile task,
pattern_17, 20 seqlets

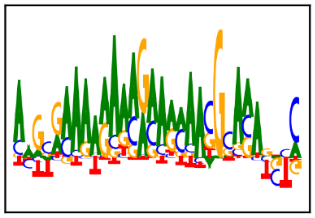

profile task,
pattern_18, 20 seqlets

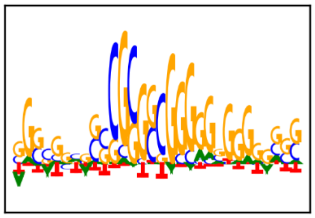

**Section 3: All baseline performances (best motif from each method)**

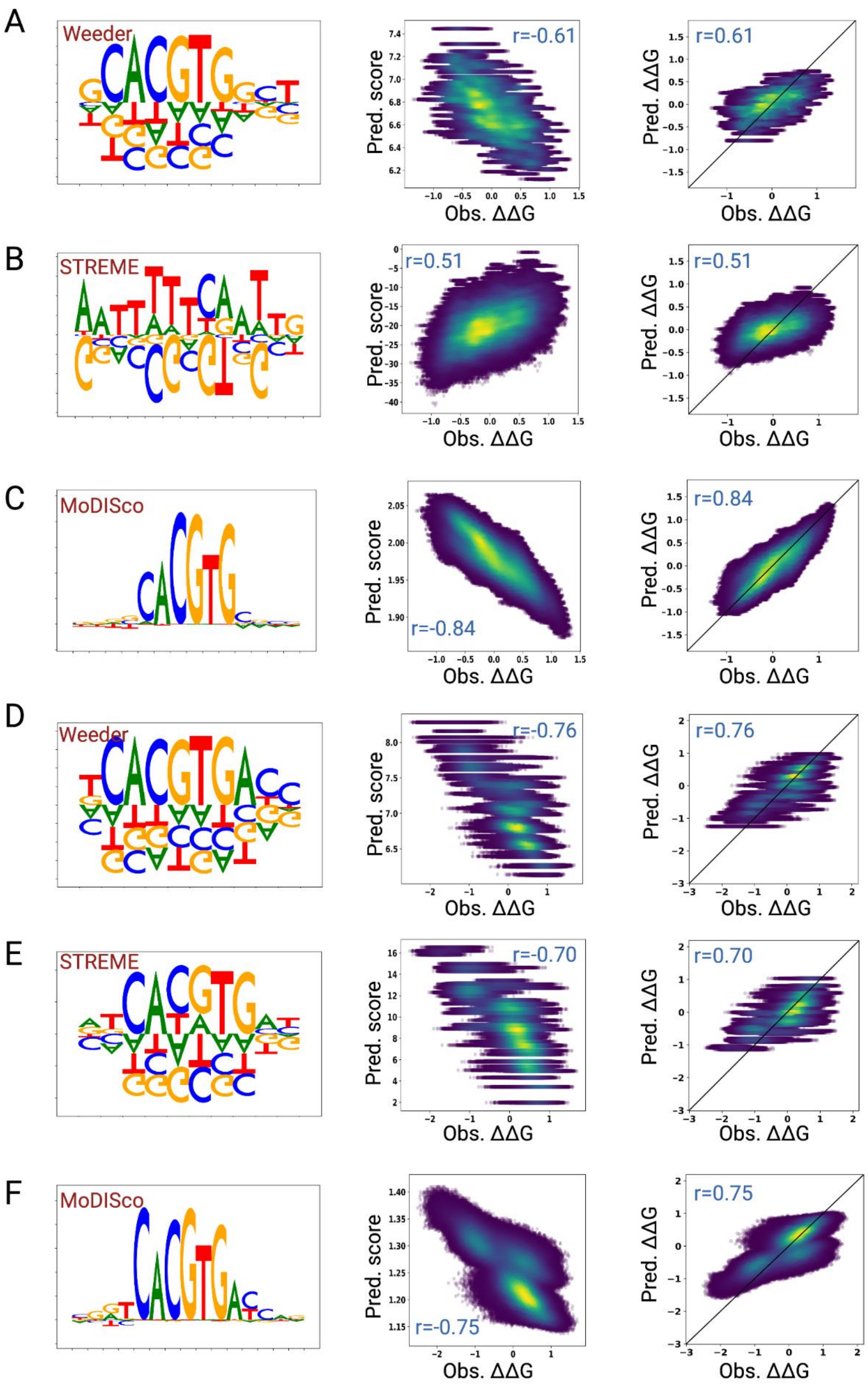


**Supplemental figure 2. Baseline evaluations for yeast PBexo models against BETseq measurements.** Best-performing (lowest RMSE) motif derived from PBexo peaks (5) (left), comparison between predicted binding scores and BET-seq-measured ΔΔGs (2) (middle), and comparison between post-calibration predicted affinities and BET-seq-measured ΔΔGs (right) for Weeder, STREME, and MoDISco prediction for Pho4 **(A-C)** and Cbf1 **(D-F)**.

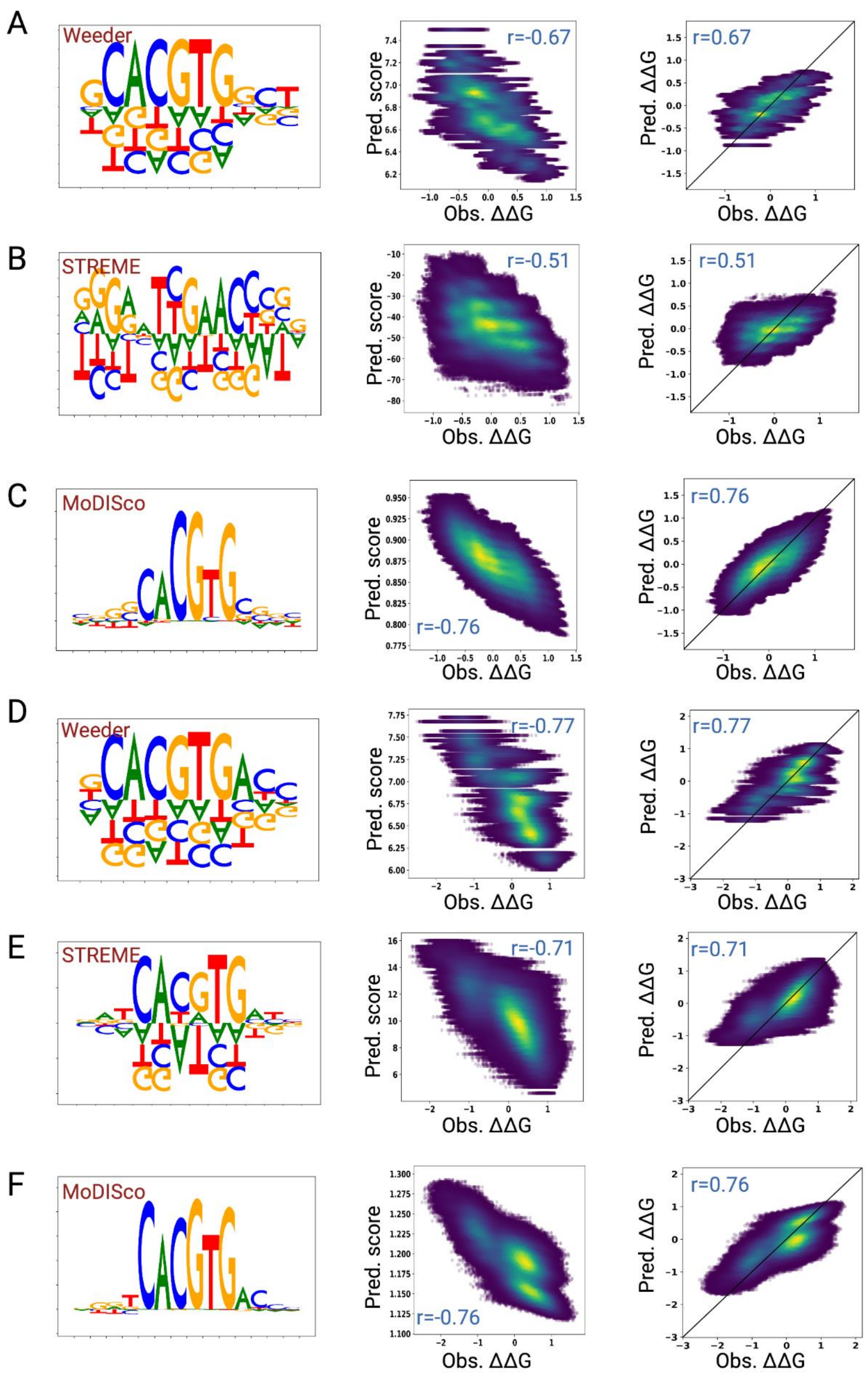


**Supplemental figure 3. Baseline evaluations for yeast ChIP nexus models against BETseq measurements.** Best-performing (lowest RMSE) motif derived from ChIP nexus peaks (left), comparison between predicted binding scores and BET-seq-measured ΔΔGs (middle), and comparison between post-calibration predicted affinities and BET-seq-measured ΔΔGs (right) for Weeder, STREME, and MoDISco prediction for Pho4 **(A-C)** and Cbf1 **(D-F)**.

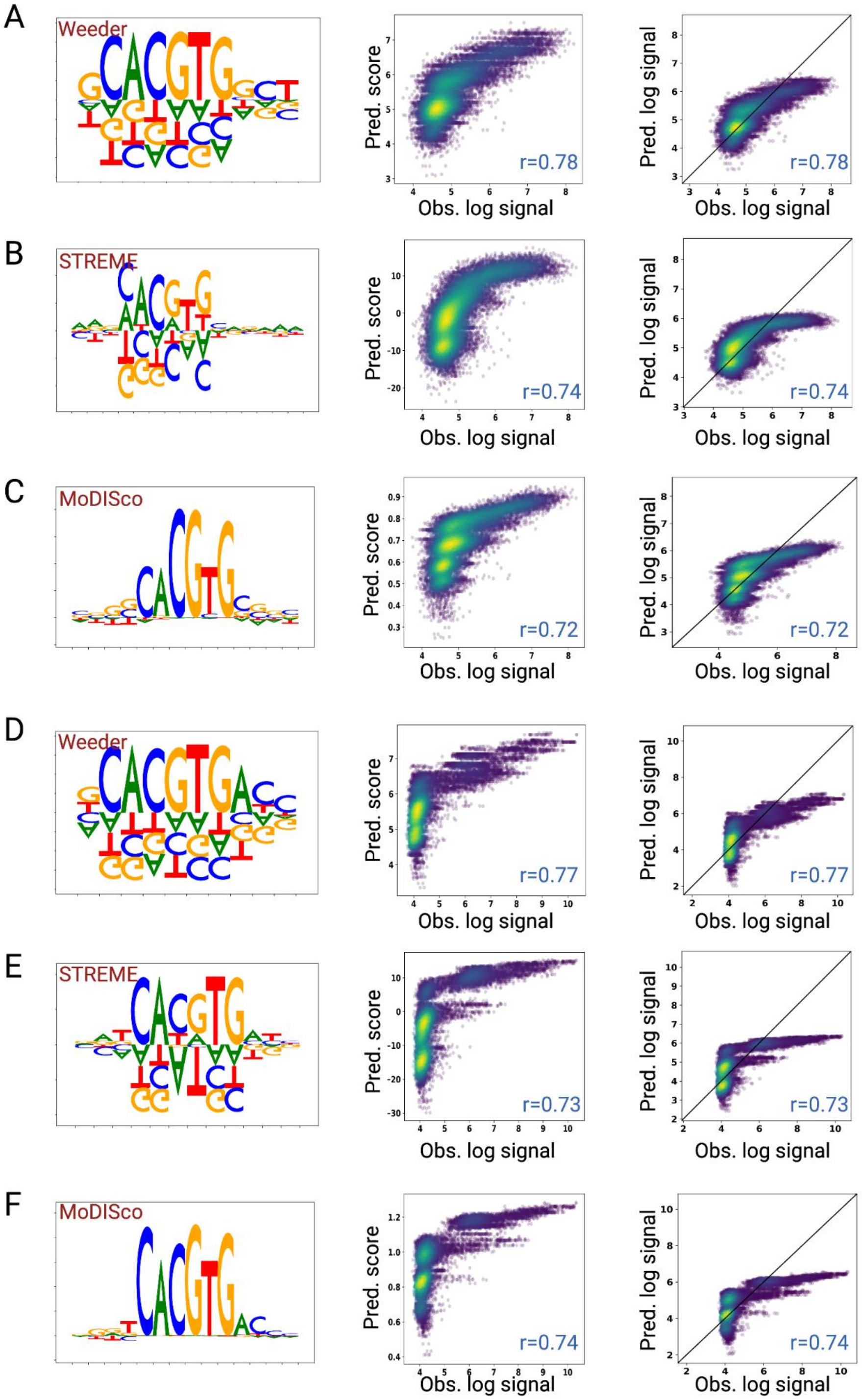


**Supplemental figure 4. Baseline evaluations for yeast ChIP nexus models against gcPBM measurements.** Best-performing (lowest RMSE) motif derived from ChIP nexus peaks (left), comparison between predicted binding scores and gcPBM log-transformed intensities (6) (middle), and comparison between post-calibration predicted affinities and gcPBM log-transformed intensities (right) for Weeder, STREME, and MoDISco prediction for Pho4 **(A-C)** and Cbf1 **(D-F)**.

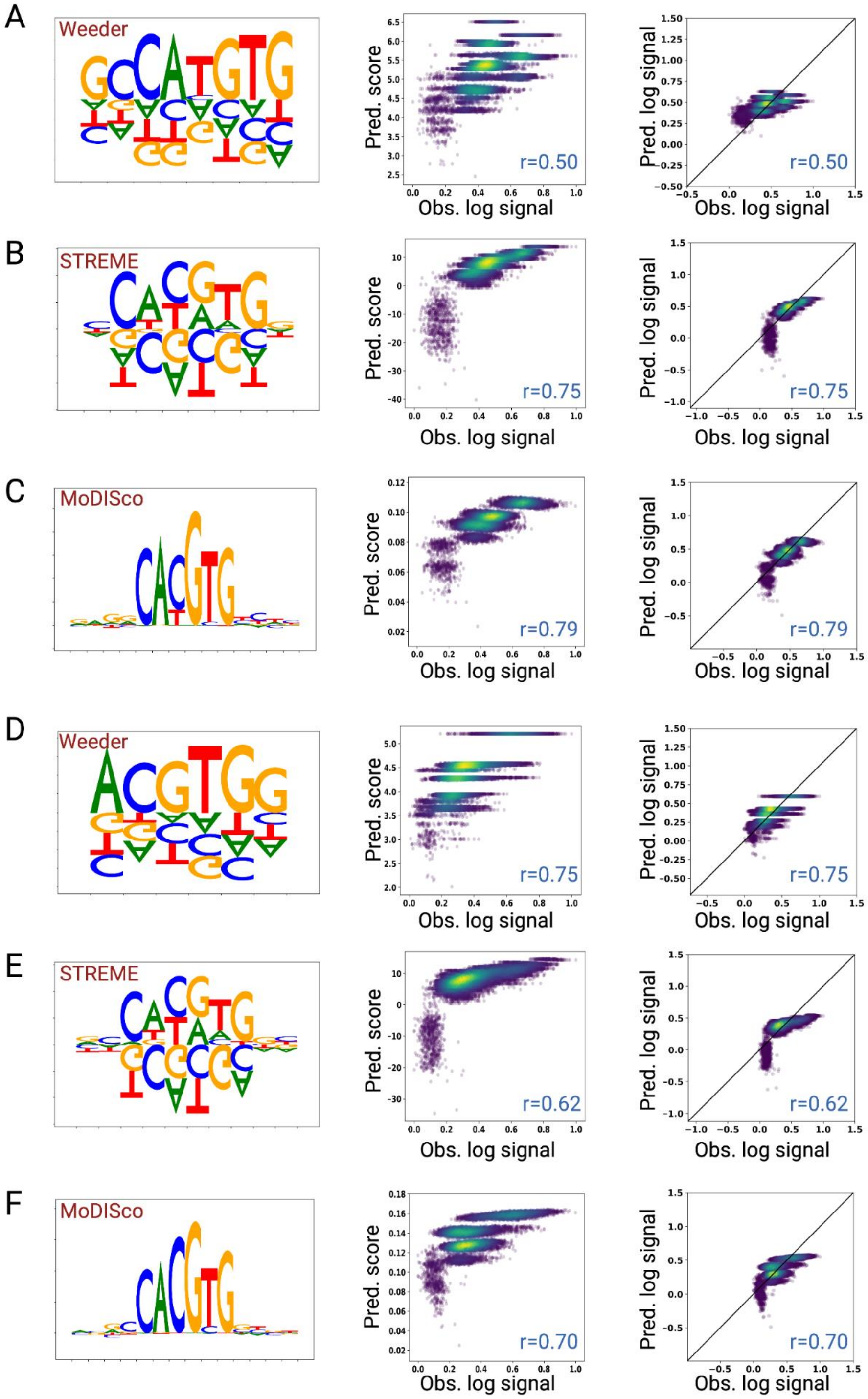


**Supplemental figure 5. Baseline evaluations for human ChIP seq models against gcPBM measurements.** Best-performing (lowest RMSE) motif derived from ChIP seq peaks in HeLa-S3 cells (left), comparison between predicted binding scores and gcPBM log-transformed intensities (7) (middle), and comparison between post-calibration predicted affinities and gcPBM log-transformed intensities (right) for Weeder, STREME, and MoDISco prediction for MAX **(A-C)** and Myc **(D-F)**.

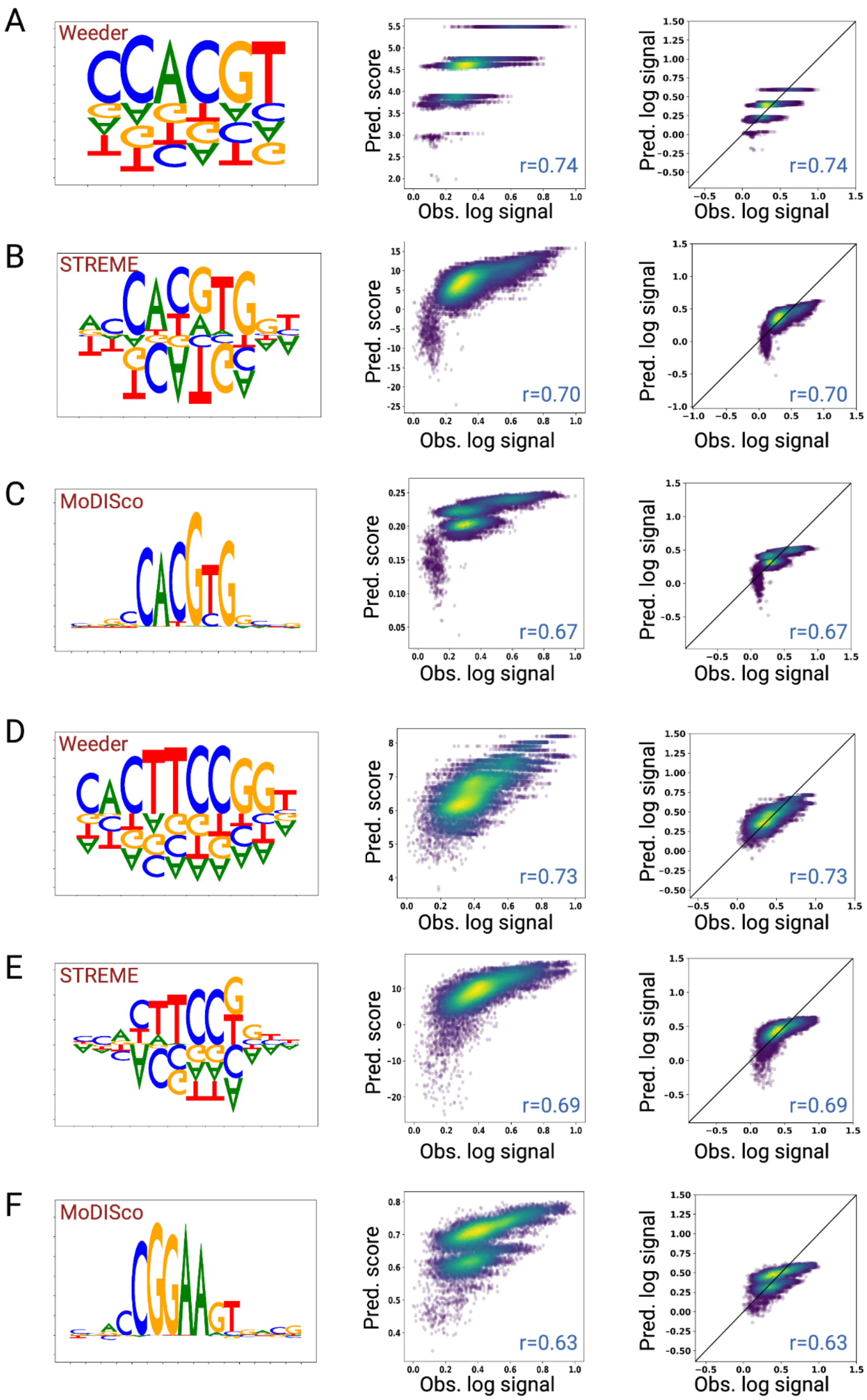


**Supplemental figure 6. Baseline evaluations for human ChIP seq models against gcPBM measurements.** Best-performing (lowest RMSE) motif derived from ChIP seq peaks (left), comparison between predicted binding scores and gcPBM log-transformed intensities (middle), and comparison between post-calibration predicted affinities and gcPBM log-transformed intensities (right) for Weeder, STREME, and MoDISco prediction for Myc in K562 cells **(A-C)** and GABPA in HepG2 cells **(D-F)**.

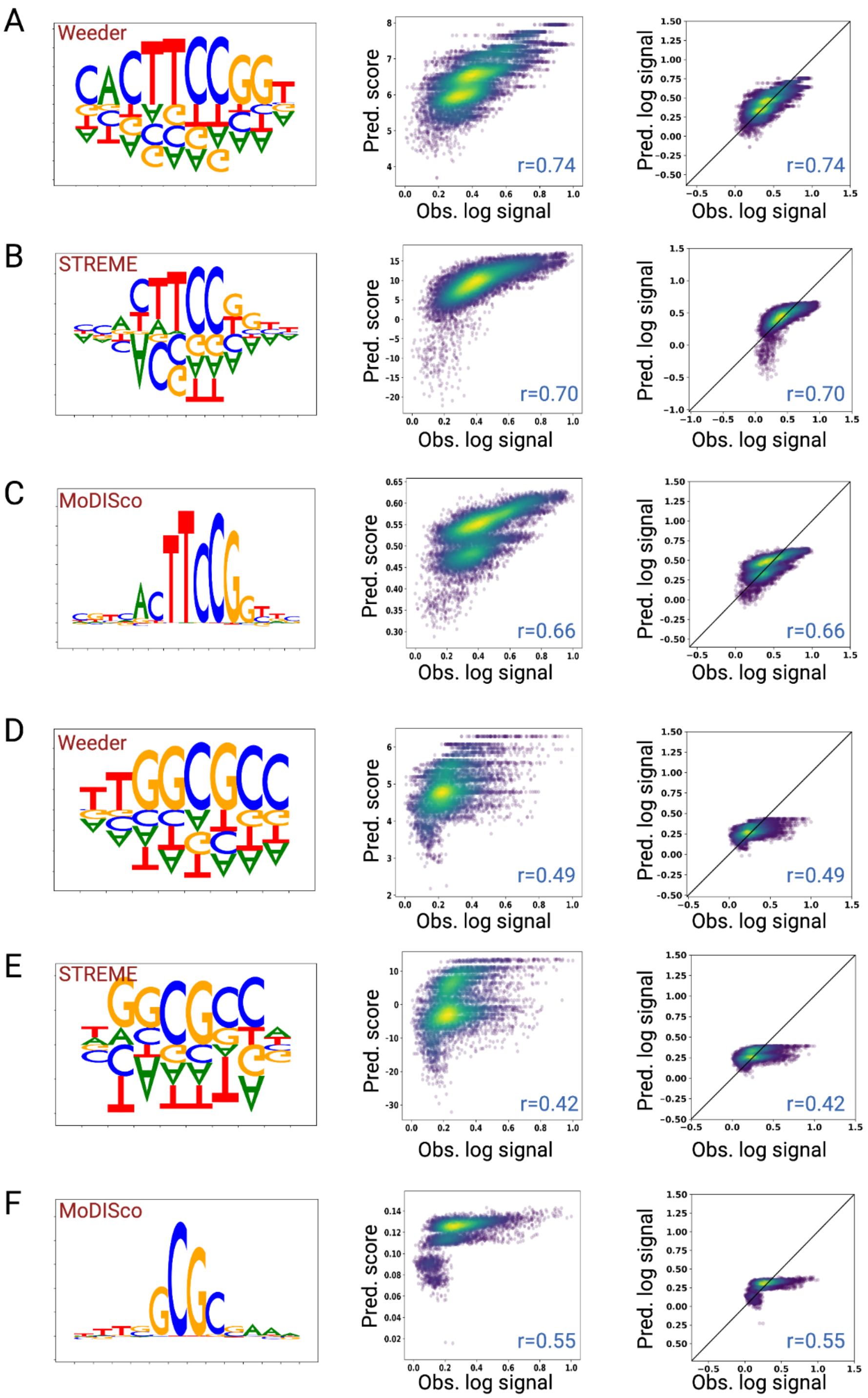


**Supplemental figure 7. Baseline evaluations for human ChIP seq models against gcPBM measurements.** Best-performing (lowest RMSE) motif derived from ChIP seq peaks (left), comparison between predicted binding scores and gcPBM log-transformed intensities (middle), and comparison between post-calibration predicted affinities and gcPBM log-transformed intensities (right) for Weeder, STREME, and MoDISco prediction for GABPA in liver cells **(A-C)** and E2F1 in MCF7 cells **(D-F)**.

**Section 4: ChIP-nexus read distribution and footprints**

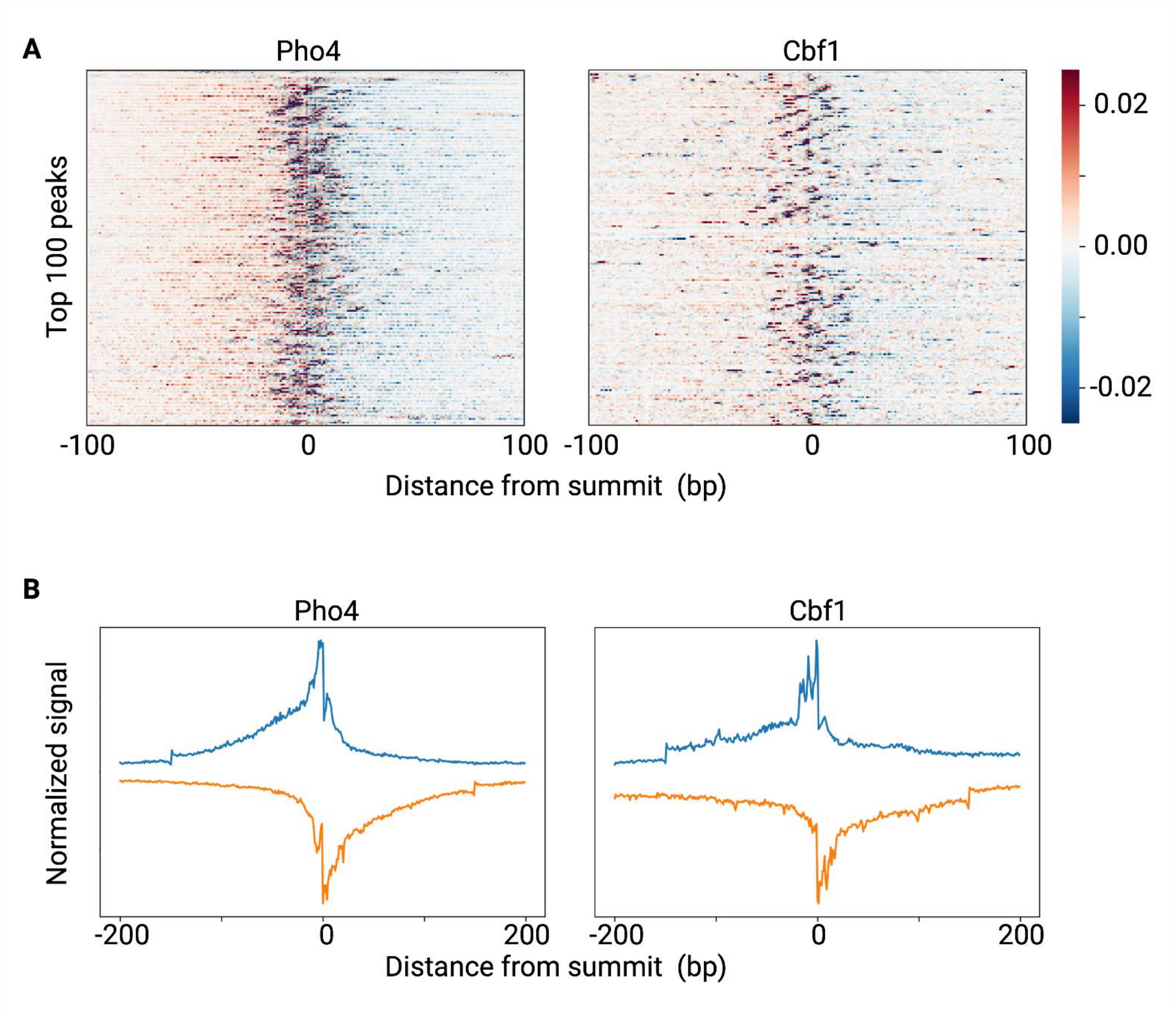


**Supplemental figure 8. ChIP-nexus read distribution and footprints.** **(A)** Profile heatmaps of Pho4 and Cbf1 ChIP–nexus data at the 100 peaks with the most ChIP–nexus reads (color depth for each strand represents normalized signal intensity). **(B)** Average Pho4 and Cbf1 ChIP–nexus footprints at the peaks.

**Section 5:  *De novo* performance of Affinity Distillation against gcPBM measurements for human TFs**

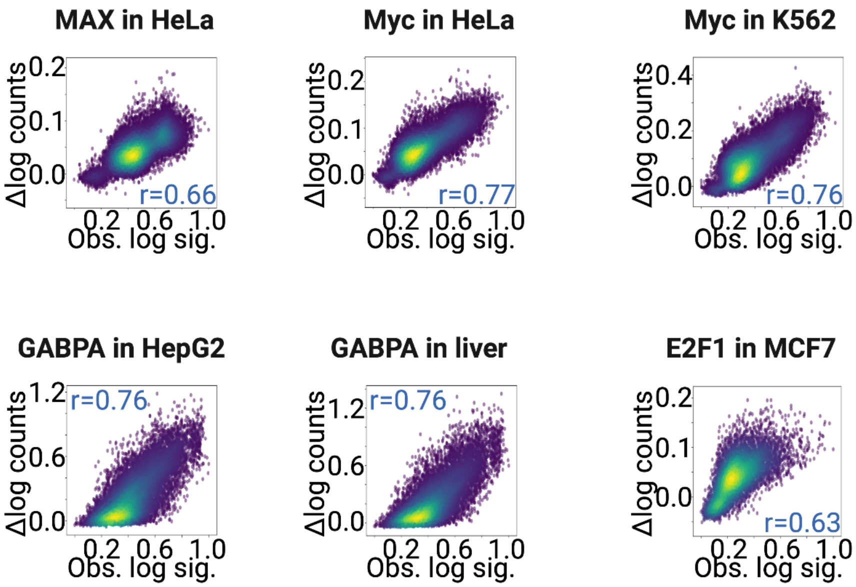


**Supplemental figure 9. Evaluation results for human ChIP seq models against gcPBM measurements.** Each panel shows the *de novo* performance (marginalization scores) against the log transformed signal intensities from the gcPBM experiments.

### Section 6: Bias correction improves Affinity Distillation

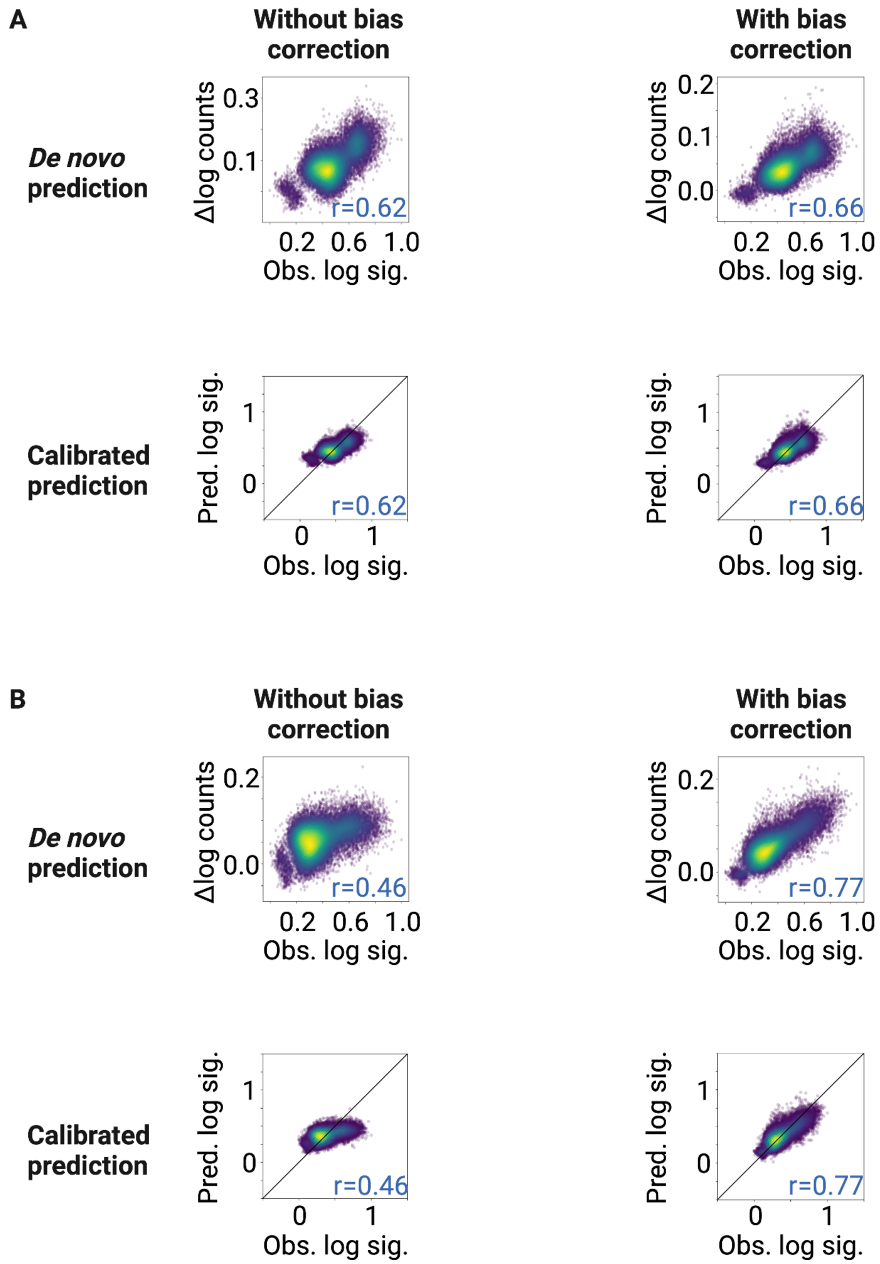


**Supplemental figure 10. Evaluation of Affinity Distillation with and without bias correction using MAX and Myc gcPBM measurements. (A)** shows the *de novo* and calibrated prediction performance of Affinity Distillation of MAX ChIP-seq models with and without bias correction in HeLa-S3 cell line. **(B)** same as A for Myc in HeLa-S3 cells.

### Section 7: Motif scoring code

```
complement = {'A': 'T', 'C': 'G', 'G': 'C', 'T': 'A'}
def getRevComp(seq):  # reverse complement function
    ret = ""
    for bp in seq.upper(): ret += complement[bp]
    return ret[::-1]

def generate_matrix(seq):
    seq_matrix = np.zeros((len(seq), 4))
    for j in range(len(seq)):
        if seq[j] == 'A':
            seq_matrix[j,0] = 1
        elif seq[j] == 'C':
            seq_matrix[j,1] = 1
        elif seq[j] == 'G':
            seq_matrix[j,2] = 1
        elif seq[j] == 'T':
            seq_matrix[j,3] = 1
    return seq_matrix

def get_PWM_score(sequence, score_matrix):
    score_len = score_matrix.shape[0]
    scores = []
    for j in range(len(sequence) - score_len + 1):
        seq_matrix = generate_matrix(sequence[j:j+score_len])
        scores.append(np.sum(score_matrix * seq_matrix))
    rc_sequence = getRevComp(sequence)
    for j in range(len(rc_sequence) - score_len + 1):
        seq_matrix = generate_matrix(rc_sequence[j:j+score_len])
        scores.append(np.sum(score_matrix * seq_matrix))
    return max(scores)

def get_PSAM_score(sequence, score_matrix):
    score_len = score_matrix.shape[0]
    score = 0
    for j in range(len(sequence) - score_len + 1):
        seq_matrix = generate_matrix(sequence[j:j+score_len])
        prod_matrix = score_matrix * seq_matrix
        score += np.prod(prod_matrix[np.nonzero(prod_matrix)])
    rc_sequence = getRevComp(sequence)
    rc_score = 0
    for j in range(len(rc_sequence) - score_len + 1):
        seq_matrix = generate_matrix(rc_sequence[j:j+score_len])
        prod_matrix = score_matrix * seq_matrix
        rc_score += np.prod(prod_matrix[np.nonzero(prod_matrix)])
    return max(score, rc_score)
```
